# Noise and determinism in Trinidadian guppy population dynamics

**DOI:** 10.64898/2026.05.17.725717

**Authors:** Harman Jaggi, Ronald Bassar, Joe Travis, Arshed Nabeel, David Reznick, Simon Levin

## Abstract

Natural populations are often nonlinear and exhibit substantial variability. A central question is how stochasticity interacts with density-dependent regulation to shape population stability. We address this using four long-term time series of Trinidadian guppies and find that their dynamics are well described by a stochastic logistic model with multiplicative environmental noise. The model predicts that stochasticity does not merely add fluctuations around deterministic carrying capacity, but alters the equilibrium structure. Using stochastic bifurcation theory, we show that increasing noise shifts the most-probable population size below the deterministic equilibrium and can push populations closer to a noise-induced bifurcation, even when mean growth rates remain positive. The effects of stochasticity across populations align with known ecological differences among streams, particularly the effects of light level and seasonality. The analysis also identifies populations most sensitive to perturbations, which are not detected by standard early warning indicators. Temporal and spectral analyses further show that intrinsic growth rate governs local recovery, while seasonal variation interacts with density-dependence to shape longer-term population fluctuations. Together, our results show that stochasticity can alter resilience and vulnerability by reshaping ecological stability landscapes.

## 1 Main text

A central goal in population ecology is to understand the causes and consequences of population fluctuations. Such variability can alter population dynamics by shifting equilibria [1], amplifying fluctuations [2], and in some cases pushing populations toward collapse [3, 4]. Ecological theory has shown that fluctuations can be caused by certain types of strong density-dependent regulation [5, 6, 7], environmental stochasticity [8, 9], demographic stochasticity [10, 11], and/or large-scale climate forcing [12, 13]. From an evolutionary perspective, population fluctuations can alter what fitness metrics are maximized under density-dependent selection [14].

The relative importance of these factors has sustained long-standing debates on the role of intrinsic regulation versus extrinsic environmental variation in determining population fluctuations [15, 16, 17]. While our understanding of the joint role of deterministic and stochastic processes in determining population variation and evolutionary trajectories has progressed, our ability to tease apart their effects and quantify the relative contributions of each to natural populations remains elusive [18, 19]. As a result, it is challenging to determine how much observed variation reflects intrinsic regulation versus stochastic fluctuations.

Stochastic differential equations (SDEs) provide a natural way to analyze both deterministic feedbacks and stochastic forces [20, 21]. While deterministic models characterize population dynamics in terms of fixed points or equilibria, stochastic dynamics are described by probability distributions of population abundance [22, 23]. In stochastic systems, long-run behavior is governed by the stationary distribution, whose most-probable state need not coincide with the deterministic equilibrium. Theory suggests that stochasticity can reshape system dynamics in non-trivial ways, and in some cases can eliminate stable states altogether [24, 25, 26, 27, 28]. Such noise-induced changes in stability can be formalized as stochastic bifurcations [29, 30]. Despite theoretical advances, empirical evidence of stochastic shifts in equilibrium structure and their consequences for population stability remain limited.

To address this gap, we develop a framework that combines data-driven model discovery with analyses of stochastic population dynamics to characterize equilibrium structure and proximity to stochastic bifurcation. Recent advances in equation discovery enable stochastic differential equations to be inferred directly from ecological time series [31, 32, 33]. Building on this approach, we derive the stationary distribution of the inferred stochastic dynamics and quantify its most-probable state. We examine how the state varies with increase in stochasticity and thereby identify shifts in equilibrium structure. These diagnostics allow us to detect proximity to noise-induced bifurcation and identify populations most sensitive to perturbations. We compute several commonly used early warning signal (EWS) metrics [34] and find such transitions are not detected by these EWS indicators based on deterministic critical slowing down [35, 36, 37].

Long-term population time series reveal temporal stability through their autocorrelation structure, which quantifies how rapidly populations recover from perturbations [38, 39, 40] and links demographic processes to fluctuation timescales. However, temporal correlations alone do not distinguish between intrinsic recovery dynamics and externally driven variability. Therefore, we use spectral analysis to examine how variance is distributed across temporal frequencies and identify signatures of environmental variability [41, 16, 42]. This allows us to separate intrinsic demographic timescales from environmental or seasonal fluctuations and assess how population dynamics filter external variability into population-level responses. Together, our approach links local recovery, long-term equilibrium behavior, and fluctuation timescales into a unified framework for analyzing stochastic population dynamics.

We apply this framework to long-term time series of Trinidadian guppies (*Poecilia reticulata*) from four experimental populations in the Northern Range mountains of Trinidad (monthly data from 2009-2025). These populations occupy two contrasting environments: two streams with intact canopy cover (Lower Lalaja and Caigual, hereafter LL and CA) and two with thinned canopies (Upper Lalaja and Taylor, hereafter UL and TY). These four populations were initiated from fish collected from the same ancestral population. The streams with thinned canopies experience higher light levels and, consequently, higher rates of primary productivity [43, 44] and higher abundances of guppy food resources like algae and invertebrates [45]. The differences among stream populations in their demographic responses to these effects offer a replicated system to examine how ecological context shapes population dynamics [46, 47, 48].

Using these long-term time series, we show that environmental stochasticity reshapes equilibrium structure, shifting the most-probable population size and driving population toward noise-induced bifurcation. Despite similar forms of density-dependent regulation, populations differ substantially in their temporal dynamics, the structure of their fluctuations, and proximity to stochastic instability, reflecting how intrinsic dynamics and environmental variability interact to shape observed dynamics. Our results provide a mechanistic link between stochastic population theory and empirical time series, highlighting the need to move beyond deterministic frameworks to account for temporally structured environmental variation.

## 2 Results

### 2.1 Guppy population densities are well-described by a stochastic logistic dynamics with state-dependent noise

Using data-driven equation discovery (see [33] and Methods), we find that population dynamics of guppies in all streams (illustrated in Figure 1) are well described by a stochastic logistic model (written in Itô form):

**Figure 1:**
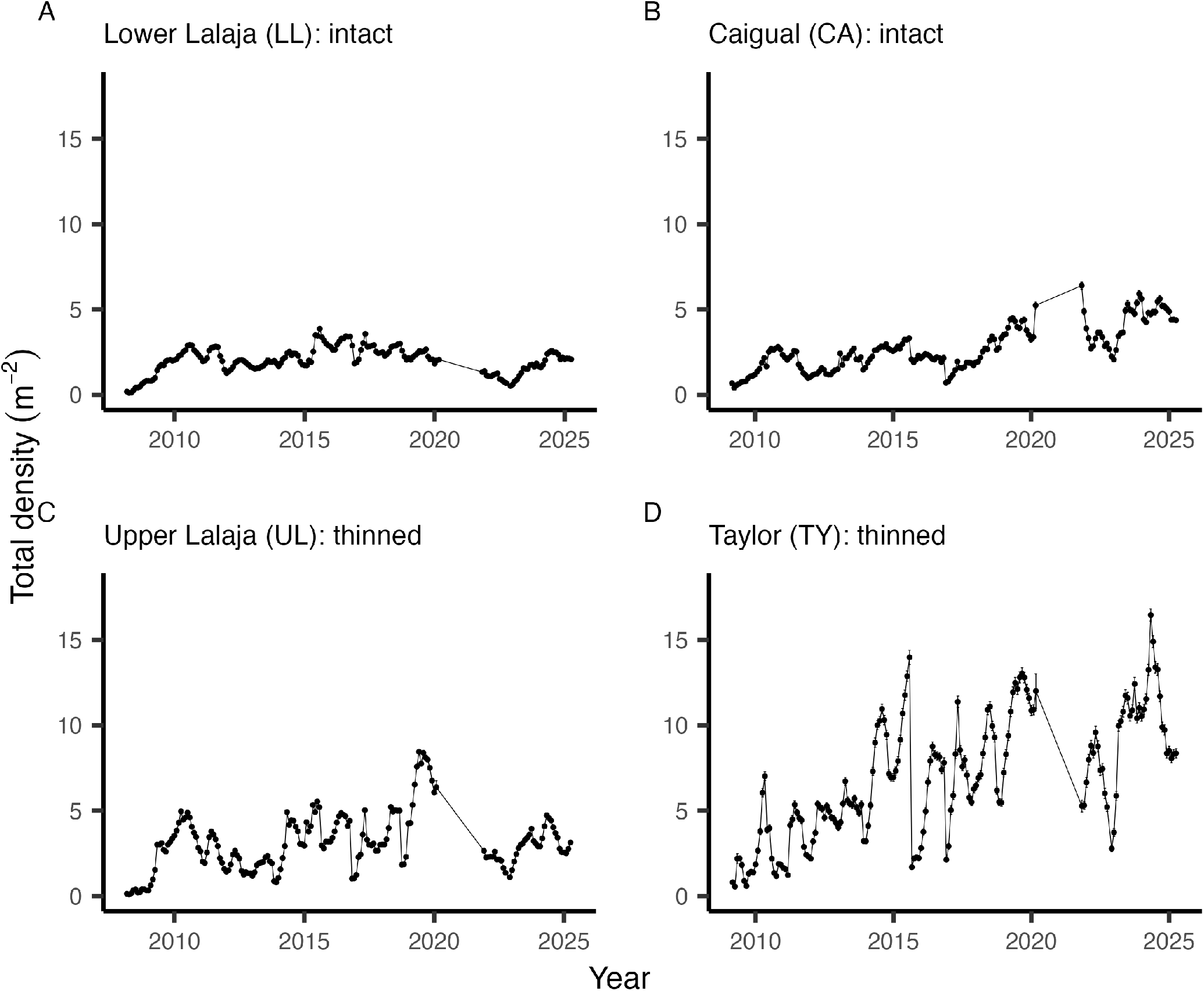
Population density of Trinidadian guppies over time. The figure plots time series of estimated total guppy density in four experimental streams: Lower Lalaja (LL), Caigual (CA), Upper Lalaja (UL), and Taylor (TY). “Intact” and “thinned” refer to canopy conditions. Points denote monthly estimates, connected by lines to show temporal trajectories, and error bars indicate uncertainty around each estimate.

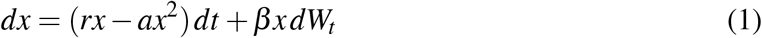

where *r* is the intrinsic population growth rate, *a* quantifies density dependence, *β* is the strength of stochasticity and *W*_*t*_ is a standard Wiener process so that *dW*_*t*_ represents Gaussian white-noise. The deterministic term, *f* (*x*) is described by a logistic growth equation, *f* (*x*) = *rx* − *ax*^2^, across all streams and is consistent with previous empirical work [48]. The stochastic component is state-dependent, with *g*(*x*) scaling with population density *x* as *g*(*x*) ∝ *x*. Thus, the magnitude of fluctuations increases with population density, consistent with multiplicative noise. Under the diffusion approximation, our results suggest environmental stochasticity as the dominant source of observed population fluctuations rather than demographic stochasticity. We do not find support for an additive, state-independent noise term on the density scale.

We find populations in thinned canopy streams exhibit both higher growth rates and stronger stochastic fluctuations, indicating noise may have stronger impact on equilibrium in these populations compared to intact-canopy populations (Fig 2 and Table 1). Across streams, the coefficient of stochastic variance is about 2.5 times larger in thinned streams (on average) than in intact streams (0.083 vs. 0.033, respectively), indicating higher fluctuations under canopy thinning. The deterministic carrying capacities were highest in TY (*K* = 7.81), and UL (*K* = 3.19), the thinned canopy streams, followed by CA (*K* = 2.96) and LL (*K* = 2.13), ), the intact canopy streams.

**Table 1:**
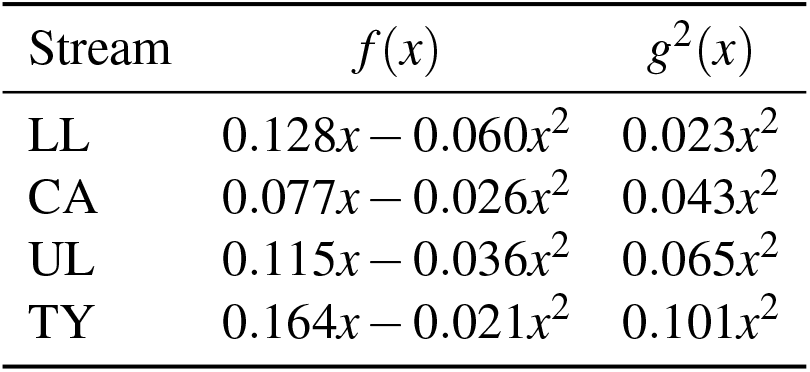
Inferred deterministic *f* (*x*) and stochastic *g*^2^(*x*) functions for guppy population dynamics.

| Stream | $f(x)$ | $g^2(x)$ |
| --- | --- | --- |
| LL | $0.128x - 0.060x^2$ | $0.023x^2$ |
| CA | $0.077x - 0.026x^2$ | $0.043x^2$ |
| UL | $0.115x - 0.036x^2$ | $0.065x^2$ |
| TY | $0.164x - 0.021x^2$ | $0.101x^2$ |

**Figure 2:**
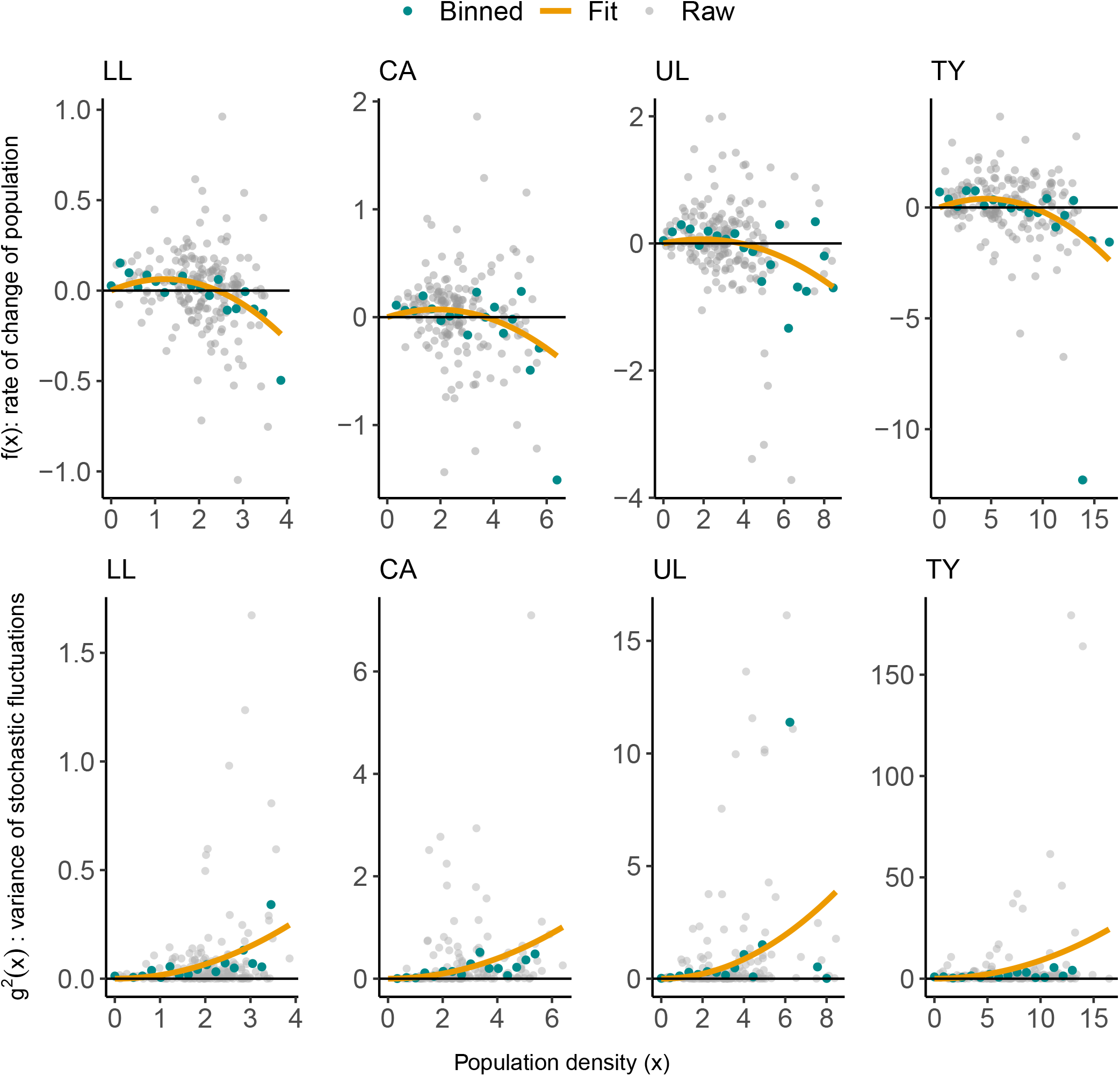
Deterministic and stochastic components of population dynamics. The figure shows deterministic *f* (*x*) and stochastic *g*^2^(*x*) component inferred from the guppy time series. The top row shows *f* (*x*) representing the rate of change in population density as a function of density. The bottom row shows *g*^2^(*x*), representing variance of fluctuations. Panels correspond to four streams (LL, CA, UL, TY), where LL, CA have intact canopy cover where UL, TY have thinned canopy cover. Grey points show raw estimates from the time series, blue points show binned averages, and orange curves show fitted functional forms.

Fig. 2 shows the inferred deterministic component, *f* (*x*) in the top row and the inferred variance of stochastic fluctuations, *g*^2^(*x*), in the bottom row. Estimated coefficients for each stream are given in Table 1. Parametric bootstrap analyses indicated that the inferred drift coefficients were consistent with logistic density dependence across streams, with positive linear terms and negative quadratic terms for all streams (Appendix Table S.1).

### 2.2 Stochasticity reshapes equilibrium structure

We find that environmental noise (*g*^2^(*x*) ∝ *x*^2^) shifts the equilibrium structure of population dynamics across all streams (Fig 3). While deterministic logistic dynamics have an equilibrium at *K* = *r*/*a*, stochastic logistic dynamics are characterized by a stationary distribution whose mode *x*^⋆^ represents the most-probable state (when such a distribution exists and is normalizable) [49]. For our stochastic logistic model with multiplicative noise, we find the stationary distribution is Gamma-distributed, with the mode given by 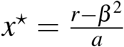. Across streams, the proportional reduction from deterministic *K* to the most-probable state *x*^⋆^ was strongest in TY (61.6%), UL (56.5%), and CA (55.9%), and weakest in LL (18.0%). Thus, increasing the strength of stochasticity (*β* ) reduces the most-probable population size and predicts the collapse of the positive equilibrium at *β* ^2^ = *r* (as shown in Fig 3). This corresponds to a *p*-bifurcation (phenomenological bifurcation) [50, 30, 51], where the positive equilibrium disappears. Such a bifurcation is understood as qualitative changes in the topology of the maxima and ridges of the stationary distribution [30]. Even when deterministic growth rate remains positive, stochasticity shifts the most-probable population size away from the equilibrium and can push populations toward instability.

**Figure 3:**
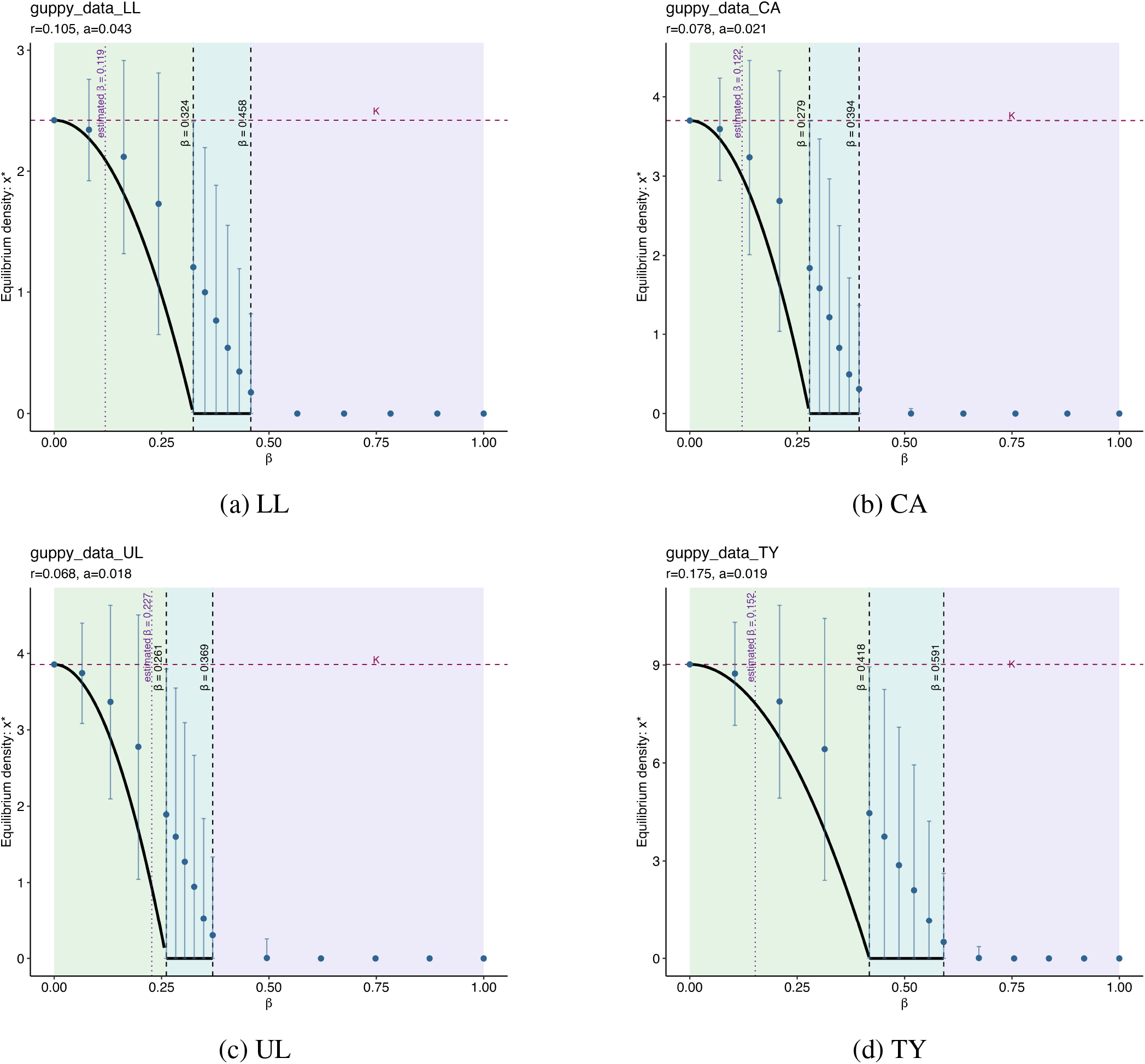
Stochastic bifurcation plot for all streams: Each panel plots the deterministic carrying capacity *K= r*/ *a* (horizontal dashed line) and the most-probable state *x*^⋆^ = (*r* − *β* ^2)^ *a* (solid black line). The vertical dashed lines at 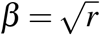 corrrespond to p-bifurcation and the second vertical dashed line at 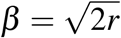 corresponds to loss of normalizability. The purple vertical dotted line denotes the empirically inferred *β* for each population. The blue dots and confidence intervals indicate the mean of the stochastic simulations.

Next, we quantify the proximity to stochastic vulnerability. We find the dynamics for all streams lie within the stable regime (*β* ^2^ < *r*), where the stationary distribution peaks at a positive abundance. However, population in the thinned-canopy stream UL lies closest to the bifurcation threshold (Fig. 3), indicating vulnerability to a noise-driven regime shift. In contrast, the estimated noise coefficient *β* for LL, CA, and TY lies within this boundary and exhibit more stable dynamical structure. Thus, we identify the population closest to stochastic instability, quantifying vulnerability beyond what is captured by deterministic growth rates or perturbations alone (as we discuss in the next section). We also calculate standard early warning indicators based on critical slowing down, but these do not identify UL as vulnerable. This is because transitions here are noise-induced, which early warning signals based on deterministic loss of stability do not capture [52, 35, 53, 37]. Corresponding early warning signal (EWS) metrics for all streams are plotted and discussed in Appendix S5.

As shown in Fig 3, we also find that the mean population size (mean of the simulations) differs from the most-probable state as stochastic coefficient *β* increases. The mean of the stochastic simulations remains positive even when the mode is close to lower abundances and eventually 0. Therefore, deterministic equilibrium and average population sizes are not representative of the typical state of the system, especially under fluctuations. This is consistent with results from stochastic dynamics theory where persistence and extinction risk depend on quasi-extinction probabilities and the lower tail of the population-size distribution rather than mean abundance [54, 20, 55]. Details for stationary distribution and stochastic bifurcation are provided in Appendix S2.

### 2.3 Population recovery is broadly consistent with local stability analysis

Autocorrelation quantifies the rate at which fluctuations in population abundance decay over time, providing a measure of temporal stability. Near equilibrium, we find stochastic logistic dynamics can be approximated by an Ornstein-Uhlenbeck (OU) process, which predicts an exponential decay of autocorrelation at rate *r*. We find that the empirical autocorrelation functions (ACFs) for the four populations are consistent with this prediction, with estimated decay close to intrinsic growth rate (*r*) as shown in Fig 4. The CA stream (intact canopy) matches the OU prediction most closely indicating rapid damping of perturbations at the rate *r*. The empirical ACF in streams LL, UL and TY initially decays at *r* and then exhibits oscillation around the exponential decay. These oscillations suggest seasonal environmental variation which we examine using spectral analysis further in the next section.

**Figure 4:**
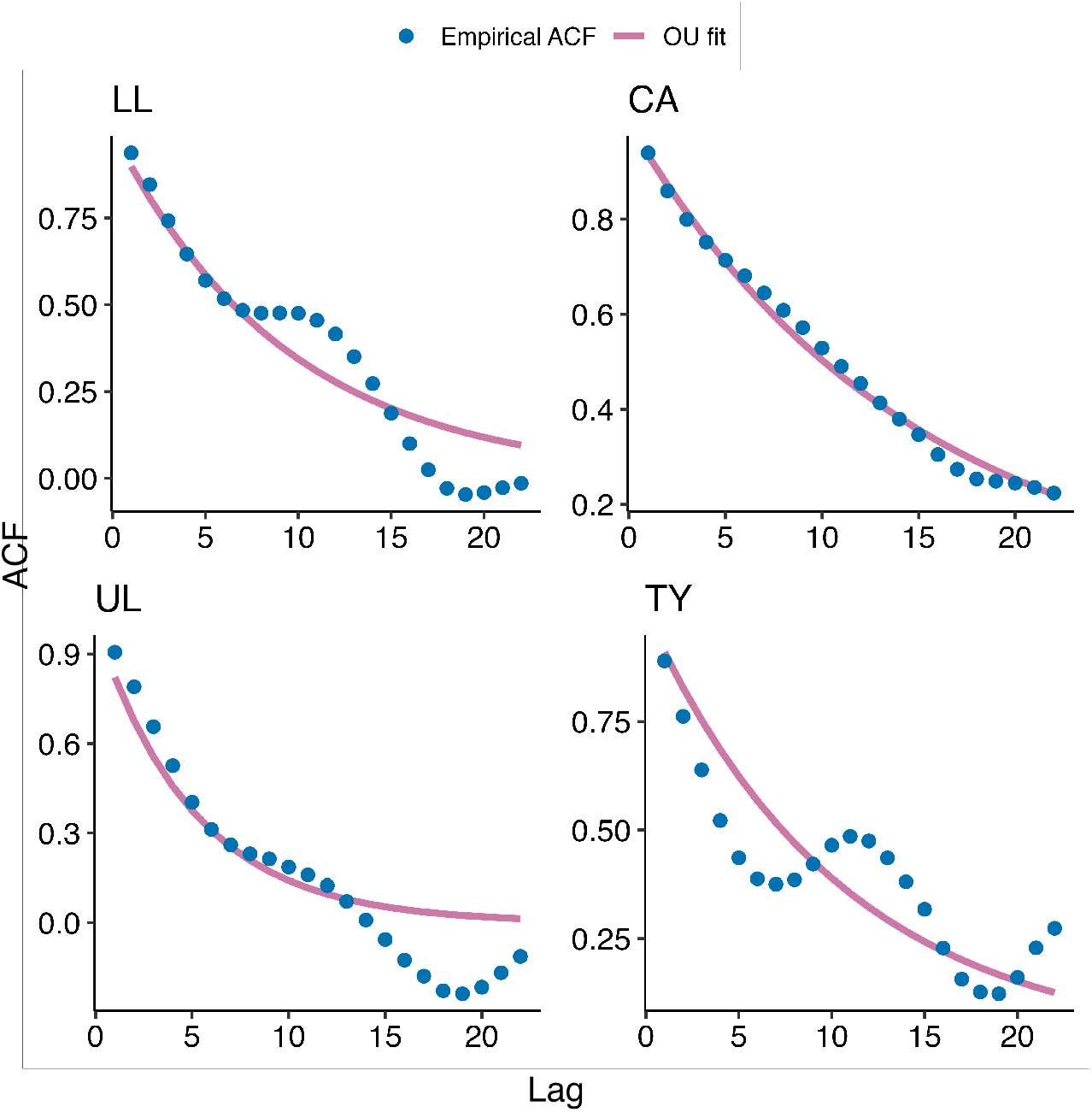
Empirical autocorrelation function (ACF) and Ornstein–Uhlenbeck (OU) fit. The blue points show empirical autocorrelation function of population density for each stream (LL, CA, UL, TY) and solid lines show the exponential decay predicted by the OU approximation near the deterministic equilibrium.

Ecologically, we find that intrinsic growth rate *r* quantifies how quickly perturbations away from *K* are damped [56, 40]. Thus, *r* sets the characteristic timescale over which fluctuations decay, implying that populations with higher *r* recover more rapidly from fluctuations. The same parameter *r* also determines proximity to stochastic bifurcation through the threshold *β* ^2^ = *r*, linking stationary distribution and recovery dynamics through a common demographic timescale.

### 2.4 Spectral analysis reveals differences in seasonal variation among populations

While autocorrelation captures the rate at which fluctuations decay, it does not tell us how the variability is distributed across temporal scales or whether fluctuations are driven by intrinsic dynamics or external variation. To examine this, we compute the power spectral density (PSD), which decomposes population variance across frequencies. We use the OU approximation to separate intrinsic dynamics from external variation by reconstructing the implied input spectrum (see Methods and Appendix S4). We find that the empirical PSD for the four streams (top panels of Fig 5) follow the Ornstein-Uhlenbeck (OU) prediction, with variance concentrated at low frequencies and decaying approximately as (*r*^2^ + *ω*^2^)^−1^ at higher frequencies, where *omega* = 2*π f* and *f* is cycles per month. This aligns with our autocorrelation results in the previous section, indicating fluctuations near equilibrium (*K*) are well described by linear dynamics.

**Figure 5:**
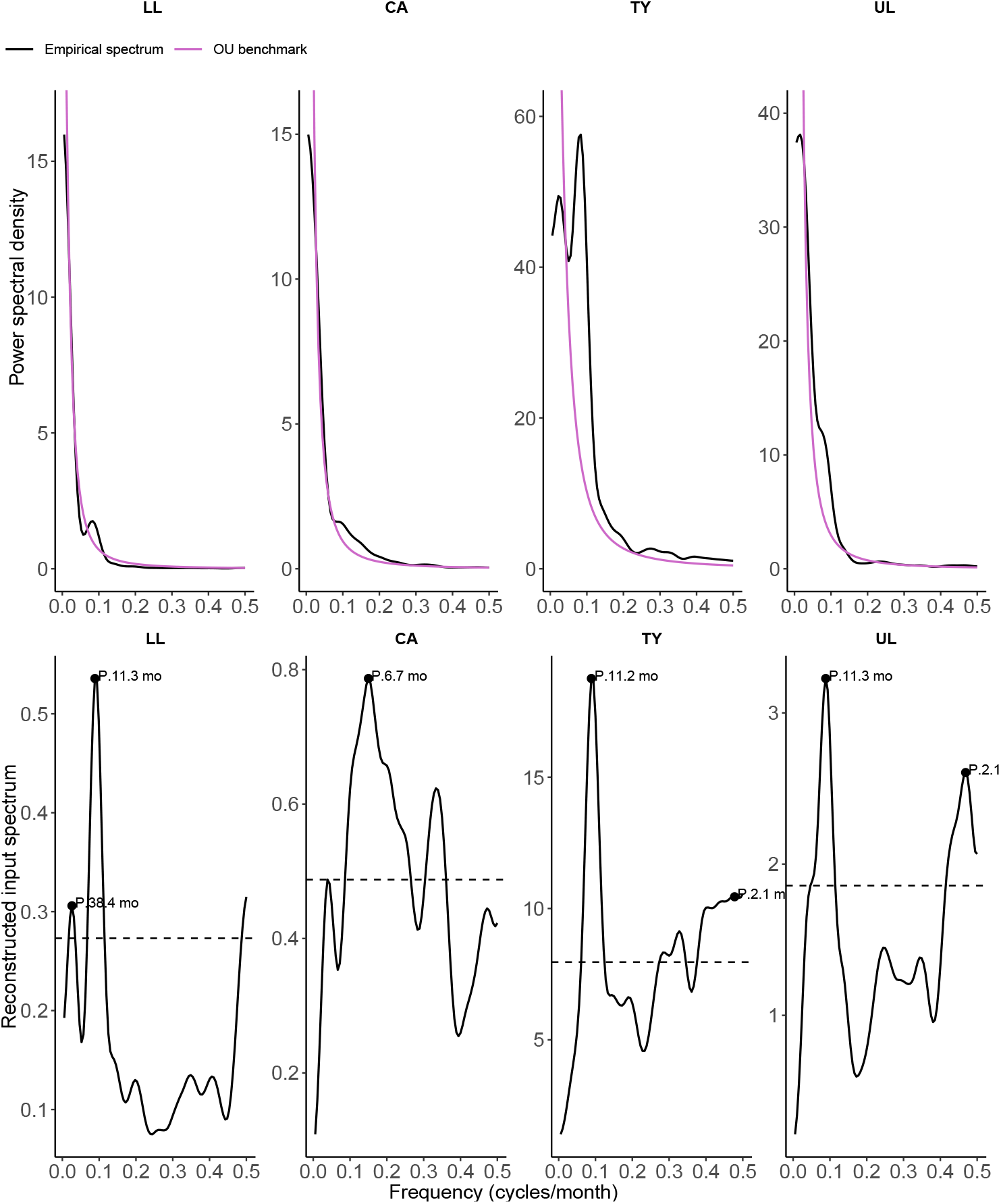
Power spectral density and reconstructed input spectrum. The top panels show the empirical power spectral density (black) and the Ornstein–Uhlenbeck (OU) benchmark (pink) for guppy populations in each stream (LL, CA, TY, UL). The bottom panels show the corresponding input spectrum reconstructed under the linear-response approximation. The dashed horizontal lines indicate the estimated white-noise reference level. The “mo” label denotes dominant peaks in the reconstructed input spectrum.

Interestingly, the reconstructed input spectrum for streams LL, UL, and TY exhibits a dominant peak at ∼11 months, indicating strong seasonal component (bottom panels of Fig 5. In contrast, intact canopy stream CA shows a much weaker seasonal signal (∼6 months) and remains closest to the OU benchmark. These peaks indicate there is a strong periodic signal with a roughly annual timescale. For a purely white noise, the observed power spectral density (*S*_*ζ*_ (*ω*)) would be flat across frequencies. Thus, departures from flatness indicate temporal structure in environmental drivers. Note that the reconstructed input spectrum should be interpreted as an effective input consistent with the linear approximation and not a unique estimate of the underlying drivers. Moreover, although all streams exhibit seasonal structure, they differ in the strength of their spectral peaks, with thinned canopy streams (UL and TY) exhibiting stronger periodic signals. Seasonal variation in mean and variance of population growth rates, linked to recruitment and survival, further supports this interpretation (see Appendix S6). The empirical mean growth rate is highest in dry season and lowest during wet season across streams.

We further computed mean-variance scaling between population density for each year and stream by fitting a log-log relationship between annual variance and mean (as shown in Appendix Fig S.6). To assess the role of seasonality, we removed the mean seasonal cycle by subtracting the long-term monthly mean (computed across years for each month) and adding back the overall annual mean. In the raw data, all streams exhibit positive mean-variance scaling, although the exponent varies across streams. After deseasonalization, this relationship breaks down in LL, UL, and TY, whereas the intact canopy stream (CA) retains the positive association (see Fig S.6). This pattern is consistent with the autocorrelation and spectral analyses, where we find CA populations dynamics are regulated by intrinsic density-dependence. However, fluctuations in other streams are more strongly shaped by external seasonal variation. Also note that TY and UL population exhibit distinct secondary peaks suggesting other sources of fluctuations and slower return rates as shown in Fig 5. Although the ecological or climatic basic of this timescale is unclear, the shared pattern between these two thinned-canopy streams may suggest a common fluctuation process. The details of the spectral analysis and reconstruction of the input spectrum are provided in Appendix S4.

## 3 Discussion

Our analyses show that environmental stochasticity can reshape the equilibrium structure of population dynamics in natural populations. In our system, the guppy population dynamics are well described by a stochastic logistic model with multiplicative environmental noise. These dynamics generate a stationary distribution whose most-probable state depends on the strength of stochasticity and density dependence. Increasing stochasticity decreases the most-probable population size (given by the mode), rather than causing populations to fluctuate around their deterministic carrying capacity. The reductions from the deterministic carrying capacity *K* to the most-probable state *x*^⋆^ were substantial across streams (ranging from 18% in LL to 61% in TY), indicating that stochasticity can strongly influence the realized population sizes. This shift is substantial even when mean growth rates remain positive and drives populations toward a stochastic bifurcation (*p*-bifurcation). Our results align with [57, 58, 14] showing that environmental stochasticity reduces expected population abundance below carrying capacity. However, the mode of the distribution provides a measure of stochastic vulnerability, as it decreases twice as fast with increasing noise and reaches 0 at *β* ^2^ = *r*, while the mean remains positive until *β* ^2^ = 2*r*.

Among the four streams, we find the population in UL stream lies closest to the *p*− bifurcation threshold, indicating increased vulnerability to fluctuations. Despite all streams being close enough to experience similar rainfall, the effect of wet season storms differs among streams. Storms in the wet season sweep guppy food resources from the streams without regard to guppy density [45]. However, the higher densities in the open canopy streams induce higher mortality rates through the lower level of per capita food after floods. Our analysis does not determine which local climatic or geomorphological conditions are most important, but it is striking that heterogeneity at small spatial scales can generate substantial differences in population dynamics. Thus, stability and resilience cannot be inferred from deterministic growth rates alone and depend on how stochasticity interacts with abiotic factors and density-dependent feedbacks to shape the long-run distribution of population sizes. This has consequences for extinction risk [3, 59], coexistence via invasion growth [60, 61] and evolutionary ecology [57, 14].

While the stationary distribution characterizes long-run population structure, temporal dynamics and linear stability analysis tell us how populations respond to fluctuations. Using local stability analysis, we derive the autocorrelation function (ACF) and predict that the rate of decay (damping rate) of fluctuations is given by the intrinsic growth rate *r*, linking demographic processes to temporal stability. However, LL, UL, and TY exhibit oscillatory deviations from exponential decay in their ACF, indicating timescales beyond local density regulation. We use spectral analysis to show that one of the reasons for these deviations is periodic seasonal variation (at approximately 11 months) in all streams except CA. Although seasonality is ubiquitous in natural systems, most population models treat environmental variation as either constant or weakly stochastic [62, 63]. Our findings highlight the role of seasonality in structuring population fluctuations and its importance in more accurately predicting system’s responses to environmental change.

To verify our findings on the role of seasonality, we computed mean-variance scaling between population density for each year and stream by fitting a log-log relationship. We then compared this to results from the deseasonalized population density. In the raw data, all streams exhibit positive mean-variance scaling. However after detrending, this relationship breaks down in LL, UL, and TY but is maintained in the intact canopy stream (CA). This pattern is consistent with the autocorrelation and spectral analyses that show CA population dynamics to be described by intrinsic density-dependent regulation. The state-space models used in Travis et al. [5] also pointed to CA as having the strongest regulation consistent with our results. LL, TY and UL are more strongly influenced by external seasonal variation.

The results across four streams highlight the importance of long-term studies with replicated populations as it helps us characterize and compare intrinsic and external factors that would otherwise be difficult to detect in single-population studies. Our analyses of long-term population time series show how local recovery from perturbations, stochastic population dynamics, and fluctuation timescales can be jointly analyzed to interpret population variability. Importantly, the inferred stochastic dynamics show that stochasticity can more than just rattle the system around equilibrium: it can reshape the effective equilibrium, stability and long-run behavior. In addition even though populations share similar logistic density dependence, they differ in their temporal dynamics such as seasonal signal, rate of fluctuation decay, and level of vulnerability. Thus, intrinsic dynamics filter environmental variability into distinct population-level responses.

Alternative data-driven approaches such as empirical dynamic modeling (EDM) focus on reconstructing nonlinear dynamics and forecasting from time series, but do not directly yield interpretable mechanistic parameters governing stability [64, 65]. Typical time-series approaches based on state-space and autoregressive models focus on parameter estimation and forecasting rather than explicitly linking dynamics to underlying mechanisms [66, 5]. In contrast, equation discovery methods allow direct inference of stochastic differential equations from data, enabling explicit links between demographic processes, stochasticity and stability properties. This provides a complementary framework for understanding population dynamics in systems where stochasticity may play a central role.

Our work also connects with early warning signals literature that has been widely used to detect regime shifts in ecological systems [67, 52]. However, these approaches are derived under deterministic bifurcation frameworks (based on critical slowing down) and may fail to detect transitions driven by stochastic effects [37]. In our system, standard early warning indicators do not identify UL population as vulnerable, consistent with the absence of a deterministic loss of stability. We also find that *r* provides a measure of resilience and sets the characteristic timescale for both bifurcation threshold (*β* ^2^ = *r*) as well as the rate at which fluctuations decay. This links growth rate to temporal stability and demography, as *r* is inversely related to generation time [56]. Thus, there is a common demographic timescale for both the stationary distribution and recovery dynamics.

Our study has several limitations. Our analysis is based on a one-dimensional representation of population dynamics, which does not explicitly account for age or stage structure, life-history tradeoffs, or species interactions [21, 68, 69, 70]. The OU approximation and spectral decomposition rely on linearization near equilibrium. While we identify the role of seasonality, we do not explicitly model the underlying environmental drivers or their mechanistic link to demographic rates [47, 71]. There is scope to further decompose intrinsic growth rate into variation from population structure and from environmental variation [72]. In addition, there is an emerging body of research that shows compelling evidence that population dynamics in microbial communities is well-explained by stochastic logistic growth [73, 74] and we find similar results for the population dynamics of Trinidadian guppies, extending their results to vertebrate populations. Our results provide a framework for integrating stochastic population theory with empirical time series to quantify how environmental variability shapes population stability. More broadly, they highlight that understanding population resilience requires moving beyond deterministic equilibria to account for how stochasticity interacts with intrinsic dynamics to shape the full distribution of population states.

## 4 Methods

### 4.1 Study system and data

We analyzed long-term population time series of Trinidadian guppies *Poecilia reticulata* from four streams in the Northern Range of Trinidad: Lalaja (LL), Caigual (CA), Upper Lalaja (UL), and Taylor (TY) (as shown in Fig. 1). All streams contain both guppies and Hart’s killifish (Anablepsoides hartii). Each stream is part of a long-term mark-recapture experiment that was initiated as part of an experimental evolution study in nature [75]. The streams range in size from 65 to 165 meters in length, are characterized by a pool/riffle structure, and contain closed populations of guppies. Two of the streams (LL and CA) have natural intact tree canopies and the other two (UL and TY) had their tree canopies thinned to increase the amount of light reaching the streams. The reduced canopy, higher light streams have higher light penetration and higher guppy population densities [76, 77, 48].

Since the beginning of the experiment in March 2008 (LL and UL) and March 2009 (CA and TY), we have returned to the streams each month to conduct mark-recapture experiments. At each capture, we removed guppies from the streams using butterfly and aquarium nets and took them to our nearby laboratory, where they were enumerated, marked, and measured for standard length and mass. Fish are individually marked with two subcutaneous, colored elastomer implants (Northwest Marine Technologies). Fish are then returned to the streams where they were captured. Further details can be found in [48].

We use the individually marked data to estimate survival, recruitment, and population size using the POPAN module of Program Mark implemented in Program R [78]. The data we present here span from March 2008 to May 2025 and include 602,836 captures of 138,932 fish. Note that the straight line between the year 2020 and 2022 reflect a break in the census because of COVID epidemic. To estimate monthly population sizes, we fitted a fully time (month) model to the individual mark-recapture data. The fully time-dependent models means that we obtained estimates of survival, recruitment, probability of capture, and population size for each stream for each month. We did not consider simpler models because our goal was to analyze the temporal trends in the data using the methods outlined above. The population-size estimates we analyze are accompanied by standard errors. These errors are in part determined by the sample sizes and probabilities of us capturing individual fish at each capture if they are alive. Our probabilities of capturing guppies if alive each month were generally between 0.80 and 0.99, averaging about 0.90, which led to estimated population sizes that were unbiased and accompanied by small standard errors. Because the standard errors are low, we chose to analyze only the estimated population sizes.

### 4.2 Stochastic differential equation discovery framework

We model guppy populations dynamics using a stochastic differential equation (SDE) that captures both the deterministic growth, *f* (*x*) (drift) and the stochastic fluctuations, *g*(*x*) (diffusion) driving the dynamics. We model population density using a one-dimensional SDE of the form:

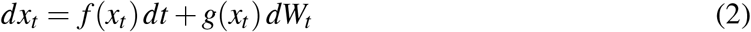

interpreted in the Itô sense. Here, *x*_*t*_ denotes population density at time *t, f* (*x*_*t*_) captures deterministic growth as a function of population density, *g*(*x*_*t*_) represents the magnitude of stochastic fluctuations around the average value, and *W*_*t*_ is a standard Wiener process (Brownian motion) [79, 80]. For the Itô SDE in eq 2, the conditional variance of short-term increments satisfies

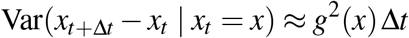

When *g*(*x*) is constant, noise strength is independent of population density and is referred to as “additive” or “state-independent” noise. When *g*(*x*) varies with *x*, noise strength depends on the current population density and is called “multiplicative” or “state-dependent” noise. In population ecology, these forms can be interpreted in relation to demographic and environmental stochasticity. Demographic stochasticity arises from randomness in individual birth and death events, whereas environmental stochasticity arises from temporal variation that affects per-capita vital rates [3, 26]. In the diffusion approximation, these contributions often appear as separate terms, with standard deviation for demographic noise scaling as 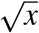 and the standard deviation for environmental noise scaling as *x*. Consequently, environmental fluctuations tend to dominate in sufficiently large populations while the demographic contribution becomes proportionally smaller [3, 26].

To infer the SDE from time-series data, we use the equation discovery framework of Nabeel et al. [33], which estimates deterministic and stochastic components directly from observed trajectories. This method proceeds in two parts: first, at each time step Δ, approximates the drift and diffusion terms using empirical estimates of the mean and variance of increments in population density. Second, sparse regression is used to identify functional forms for deterministic *f* (*x*) and stochastic *g*(*x*) parts. Lastly, to assess uncertainty in the inferred drift terms, we performed a parametric bootstrap for each stream.

### 4.3 Stationary distribution

To characterize equilibrium structure, we analyze the stationary distribution of the inferred stochastic dynamics:

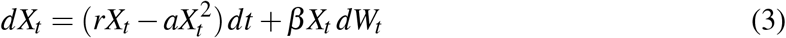

We derive the stationary probability distribution *p*^∗^(*x*) from the corresponding Fokker-Planck equation under natural boundary conditions (*x* = 0 and *x* → ∞) (see Appendix A for derivation). The resulting stationary density is a Gamma distribution of the form:

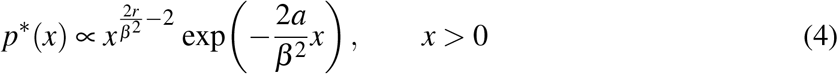

The mean and variance of the stationary population density are easily evaluated for the Gamma distribution and given by:

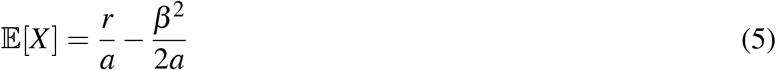

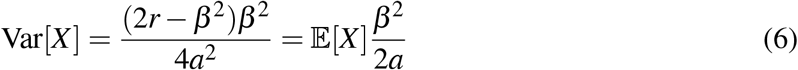

The expression is well-defined for *β* ^2^ < 2*r*, ensuring normalizability of *p*^∗^(*x*). Both expressions show that increasing stochasticity (*β* ) decreases mean population density and increases variance around equilibrium.

#### 4.3.1 Noise-induced shifts at stationarity

In deterministic systems governed by ordinary differential equations 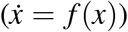, equilibrium or fixed points describe long-term steady states. For stochastic systems, an analogue is required to characterize the stationary behavior of the dynamics. In our case, the mode (*x*^⋆^) of the stationary distribution plays this role. This is because the mode represents the most probable state of the system around which probability mass accumulates and spends the largest fraction of time at stationarity [30]. While the mean 𝔼 [*X* ] captures the average abundance across all stochastic realizations, the mode identifies the density around which the population most frequently fluctuates [30, 51].

Because the stationary distribution in Eq. (4) follows a Gamma distribution with shape *α* = 2*r*/*β* ^2^ − 1 and scale *θ* = *β* ^2^/(2*a*), the mode of the Gamma distribution is easily calculated as:

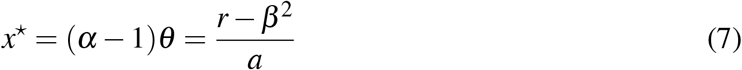

which exists for *β* ^2^ < *r*. We want to note that alternative representations based on convective field decomposition [36, 30] are provided in Appendix S2.3.

### 4.4 Local linear stability around deterministic equilibrium

We analyze the dynamics near the deterministic equilibrium 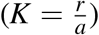 of equation 3 by linearizing the SDE around *K*. We linearize by writing *X*_*t*_ = *K* + *ε*_*t*_, where *ε*_*t*_ represents deviations from equilibrium. Around this equilibrium (*K*), the guppy population exhibits approximately linear fluctuations, described by an Ornstein-Uhlenbeck (OU) process (see Appendix for details):

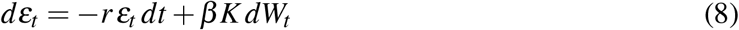

where *r* is the intrinsic population growth rate and corresponds to damping or relaxation rate and *β K* corresponds to the noise amplitude. To measure system’s stability and the timescale of decay around equilibrium, we calculate the stationary autocovariance for the OU process

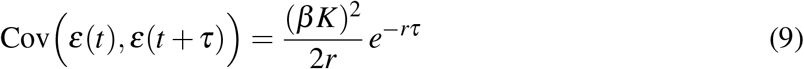

The autocorrelation function (ACF) is given by ACF_*X*_ (*τ*) ≈ *e*^−*rτ*^ . Theoretically, small perturbations away from *K* decay exponentially at the rate *r* and fluctuations have variance proportional to (*β K*)^2^.

#### 4.4.1 Fitting model to empirical autocorrelation

For each stream, we want to test if the data exhibit OU-like decay around *K*. So we first estimate the drift and diffusion coefficients *f* (*x*) = *rx* − *ax*^2^ and *g*^2^(*x*) = *β* ^2^*x*^2^ (Table 1) using Nabeel et al. [33] . From this we get the deterministic equilibrium 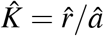 and the noise coefficient 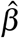. We then center population time series by defining 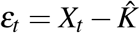 and compute the empirical autocorrelation at lag *k* using the `acf` function in `R`. Lag *k* is measured in months.

Under the OU approximation (equation S29), the autocorrelation should decay exponentially as *ρ*(*k*) = exp(−*rk*) for lag *k*. For each stream, we estimate the decay rate by fitting an exponential curve to the empirical ACF using the `nls` function. We compare the fitted decay rate 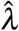 to 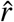 from the SDE model to check how well the OU approximation captures the observed decay of correlations. The OU model predicts a stationary variance 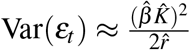 . We also compare the empirical variance of *ε*_*t*_ with the OU prediction.

### 4.5 Spectral analysis of population fluctuations

Population fluctuations can be studied in the time domain or in the frequency domain. To examine how variance is distributed across different frequencies, we consider the power spectral density (PSD). For a stationary process, the power spectral density is the Fourier transform of the autocovariance function [81, 42]. Thus, temporal stability can be characterized by the decay rate of the autocorrelation function (relaxation rate *r* in our OU approximation) or by the concentration of spectral power at low frequencies. The height of the spectrum reflects the amplitude of stochastic fluctuations, while the distribution of power across frequencies reveals how variance is partitioned between fast and slow timescales. Systems with greater temporal stability concentrate power at high frequencies (rapid recovery), whereas those near instability exhibit large low-frequency power, indicating persistent fluctuations.

Linearization of the stochastic logistic equation near its deterministic equilibrium (*K* = *r*/*a*) yields a first-order linear time-invariant system driven by a stochastic input *ζ* (*t*) = *β Kη*(*t*). In the signal processing literature, the relationship between the input (environmental) and output (population) spectra is given by:

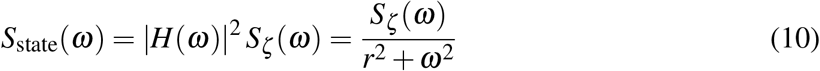

where *H*(*ω*) = 1/(*r* + *iω*) is the transfer function corresponding to the exponential impulse response *h*(*t*) = *e*^−*rt*^ (see Appendix S4 for details).

We estimate the empirical state spectrum *S*_state_(*ω*) from the observed population densities *x*_*t*_ using a smoothed periodogram implemented in the `spec.pgram` function (`R` base package). The time series was linearly detrended and mean-centered to satisfy the stationarity assumption required for spectral estimation. Frequencies ( *f* ) are expressed in cycles per month, with angular frequency *ω* = 2*π f* . The spectral density is normalized so that its integral matches the empirical variance of *x*_*t*_.

We can reconstruct the implied input spectrum using (10):

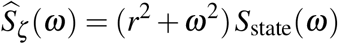

which characterizes the frequency structure of effective environmental variability under the linearized dynamics. Note that the inferred 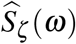 represents an effective input spectrum consistent with the observed dynamics under a linear-response approximation, rather than a uniquely identifiable environmental forcing. In addition, in applying Eq. (10), we assume that the dynamics are locally linear near equilibrium and that multiplicative noise can be approximated as additive here. Thus, the observed population variability can be interpreted as a filtered response to external fluctuations, allowing us to identify the timescales at which environmental variability influences population dynamics.

Lastly, to compare and interpret empirical state spectrum *S*_state_(*ω*) against a theoretical benchmark, we use the OU spectrum. For an Ornstein–Uhlenbeck (OU) process driven by white noise of variance *B* = (*β K*)^2^ (for our linearized stochasic logistic model), the power spectral density is given by:

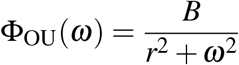

## 5 Acknowledgements

HJ thanks Stephan Munch, Shripad Tuljapurkar, and Vivek Jadhav for their helpful feedback. AN thanks Vishwesha Guttal for valuable discussions. HJ and AN thank the ICTS-ICTP Winter School (ICTS/qsb2025/01) on Quantitative Biology for engaging lectures and sessions. HJ thanks Tim Coulson for introducing to the guppy project. HJ acknowledges the support of the Princeton University Dean for Research, the High Meadows Environmental Institute and funding support from William H. Miller III.

## Data sharing

Data and code are provided as private-for-peer review via the following link: https://doi.org/10.5281/zenodo.20252187

## Appendices

### Appendix S1 Inferred coefficients table

Coefficients correspond to the fitted drift function *f* (*x*) = *rx* + *ax*^2^. We obtained the bootstrap standard errors and percentile 95% confidence intervals by simulating replicate time series from the fitted stochastic logistic model for each stream and then refitting the data-driven SDE discovery method to each simulated replicate. Standard errors were computed as the standard deviation of bootstrap estimates, and 95% confidence intervals were taken as percentile intervals from the bootstrap distribution.

**Table S.1:**
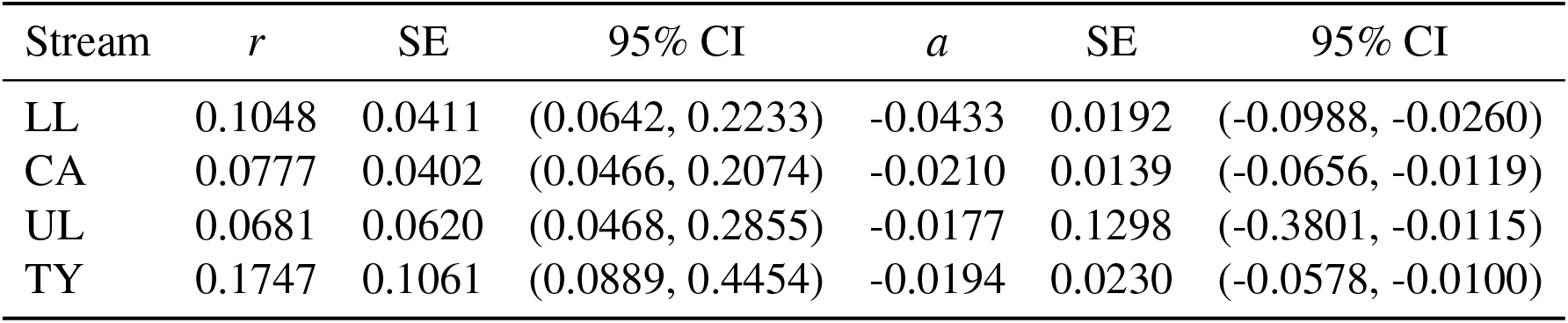
Parametric bootstrap uncertainty for the inferred drift coefficients in each stream.

| Stream | $r$ | SE | 95% CI | $a$ | SE | 95% CI |
| --- | --- | --- | --- | --- | --- | --- |
| LL | 0.1048 | 0.0411 | (0.0642, 0.2233) | -0.0433 | 0.0192 | (-0.0988, -0.0260) |
| CA | 0.0777 | 0.0402 | (0.0466, 0.2074) | -0.0210 | 0.0139 | (-0.0656, -0.0119) |
| UL | 0.0681 | 0.0620 | (0.0468, 0.2855) | -0.0177 | 0.1298 | (-0.3801, -0.0115) |
| TY | 0.1747 | 0.1061 | (0.0889, 0.4454) | -0.0194 | 0.0230 | (-0.0578, -0.0100) |

### Appendix S2 Stationary probability distribution for the SDE

From the equation discovery framework discussed in Nabeel et al. [1], we find that the guppy dynamics across streams can be modeled as a stochastic logistic model:

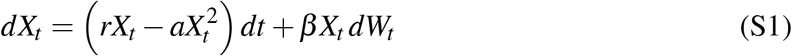

with *r, a, β* > 0. In this stochastic dynamical equation (SDE), *X*_*t*_ denotes the stochastic population size and is a random variable.

To describe the probability density of *X*_*t*_ over time, we use the Fokker-Planck equation as it governs how the distribution of population sizes evolves. Specifically, if *p*(*x, t*) is the probability density, then *p*(*x, t*) *dx* = Pr(*x* ≤ *X*_*t*_ ≤ *x* + *dx*) gives the probability of finding the population in the interval [*x, x* + *dx*] at time *t*. The deterministic part (or drift) is 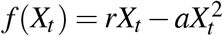, and the stochastic part (or diffusion coefficient) is 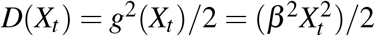.

The Fokker-Planck equation for the probability density *p*(*x, t*) of the random variable *X*_*t*_ is

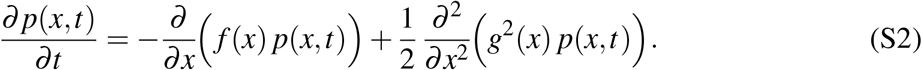

with

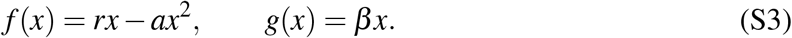

At stationarity, the probability density does not change in time and so:

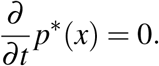

Thus, the Fokker-Planck equation reduces to a balance of the probability flux or current given by [2]:

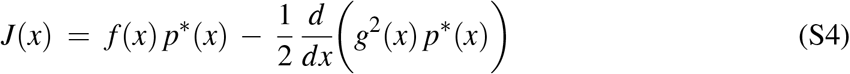

where *p*^∗^(*x*) is the stationary distribution. Note that we will return to probability current in the section on stochastic bifurcations (in Appendix S2.3). For the stochastic logistic model on (0, ∞), we assume both *x* = 0 and *x* → ∞ are natural boundaries. As discussed in Risken and Haken [3], for a stationary process, the probability flux must be constant. With natural boundary conditions at *x* = 0 and *x* → ∞, the probability flux must vanish and therefore

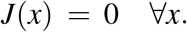

This condition ensures detailed balance, *i*.*e*, at each population size the probability mass does not flow out of the system so that the stationary distribution *p*^∗^(*x*) is conserved [3, 4]. Since *J*(*x*) = 0, we get

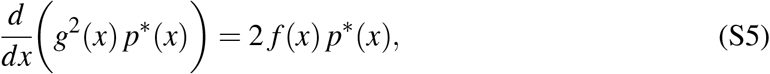

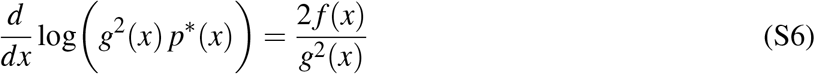

Integrating (S6) yields the standard form of the stationary density for our one–dimensional SDE:

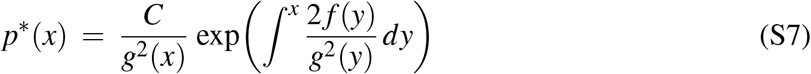

where *C* is a normalizing constant chosen such that 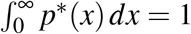. This expression is the general stationary solution for stochastic population models with a single state variable under natural boundary conditions [4].

#### S2.1 Logistic drift and multiplicative noise

We have *f* (*x*) = *rx* − *ax*^2^ and *g*^2^(*x*) = *β* ^2^*x*^2^

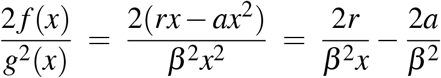

Hence

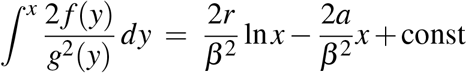

Substituting into (S7), the stationary distribution is

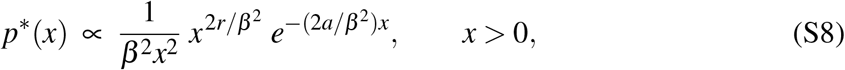

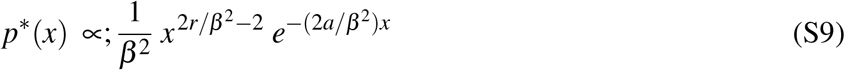

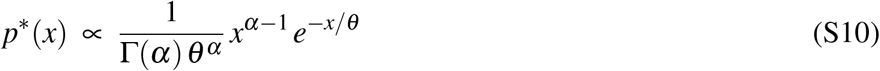

which is a Gamma distribution with shape (*α*) and scale (*θ* )

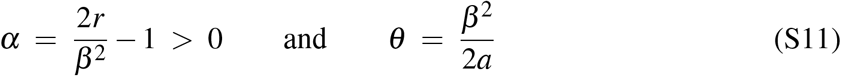

The condition *α* > 0 (2*r* > *β* ^2^) ensures that the stationary distribution does not diverge near very small population sizes (*x* → 0). Thus, the total probability remains finite and the distribution can be normalized.

#### S2.2 Stationary mean and variance

For a Gamma distribution with shape *α* and scale *θ*, the mean and variance are given by:

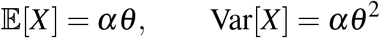

Substituting the parameters from equation (S11) we get the expected population is:

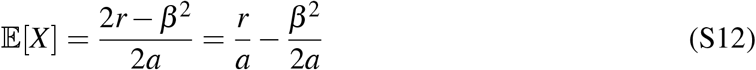

Thus in the presence of environmental stochasticity (*β* ), the expected population size 𝔼[*X* ] is strictly less than the deterministic carrying capacity 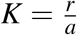, reflecting a noise-induced reduction of effective carrying capacity due to fluctuations.

And the variance is:

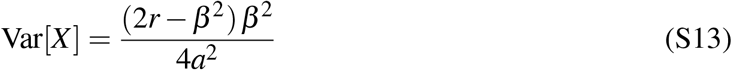

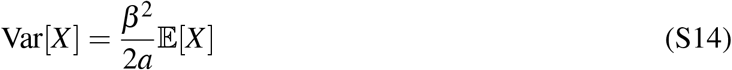

#### S2.3 Convective field representation

The stationary analysis can be equivalently expressed using the convective field formalism of Mendler et al. [2], which decomposes the SDE into deterministic drift and noise-induced drift terms as shown below:

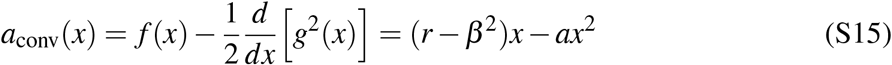

The zeros of *a*_conv_(*x*) correspond to stationary extrema of *p*^∗^(*x*). The interior root *x*^⋆^ = (*r* − *β* ^2^)/*a* coincides with the mode derived above, confirming that the deterministic growth term ( *f* ) and the noise-induced drift 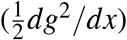 balance at this point. At *β* ^2^ = *r*, this zero collides with the boundary, producing the stochastic *p*-bifurcation. In this view, the most-probable state is not a fixed point of the deterministic system but the point where deterministic and stochastic forces exactly equal and opposite.

### Appendix S3 Linearization around carrying capacity and OU approximation

The deterministic equilibrium (or carrying capacity *K*) for the SDE in S1 is 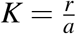 .

Let *x* = *K* + *ε*, where *ε* is the deviation from equilibrium *K*. Expanding the deterministic term *f* (*x*) = *rx* − *ax*^2^ around *K* gives

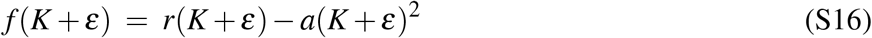

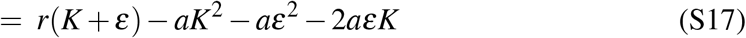

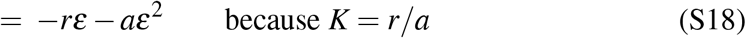

So to linear order in *ε, f* (*x*) ≈ −*rε*. For the stochastic term *g*(*x*) = *βx*, we approximate near *K* as

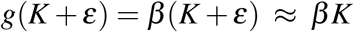

ignoring *Kε*. We also retained the first-order dependence of the stochastic term near *K* by writing *g*(*K* + *ε*) ≈ *β K* + *β ε*. In our data, incorporating *Kε* did not improve the autocorrelation fits relative to the additive-noise.

Thus, the linearized dynamics of deviations are approximated as:

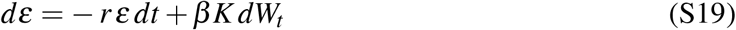

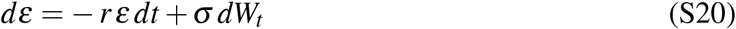

where *σ* = *β K*. Equation (S20) is an Ornstein–Uhlenbeck (OU) process that describes mean-reverting fluctuations around equilibrium. Ecologically, the OU approximation provides a local description of stochastic population dynamics near carrying capacity *K*. In this neighborhood, perturbations away from *K* decay exponentially, and the strength of fluctuations is set by the noise amplitude *β* and equilibrium size *K* as we in the section below.

#### S3.1 OU process and properties

##### S3.1.1 Mean, variance, and autocovariance

The OU process has well-known statistical properties [4, 3]. Following standard results for the process [see, e.g., 4, Chapter 5.2], the mean, variance, and autocorrelation can be derived directly from the linearized SDE:

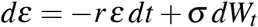

Using the integrating factor *e*^*rt*^, the solution is given by:

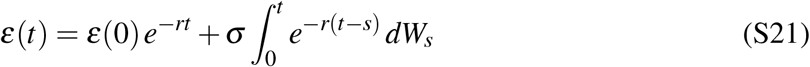

Taking expectation:

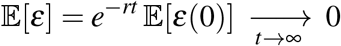

To obtain the stationary representation, we let the start time *t*_0_ → −∞ to remove dependence on initial conditions, ensuring that *ε*(*t*) has the same distribution for all *t*. In this form,

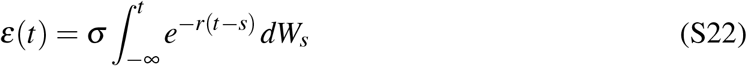

Hence, under linearization *X* = *K* + *ε*, the mean is given by: 𝔼[*X* ] ≈ *K*

The stationary variance is

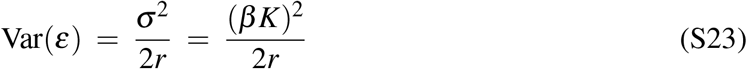

Using the stationary representation in (S22), we know: 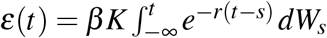

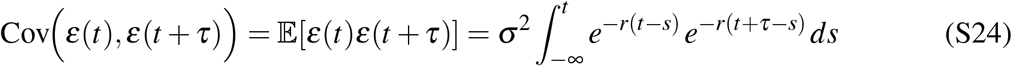

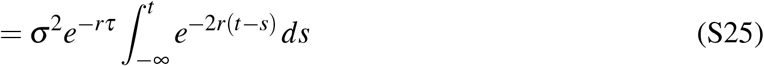

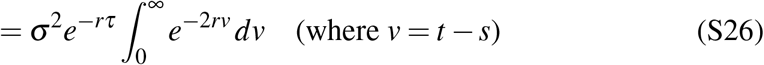

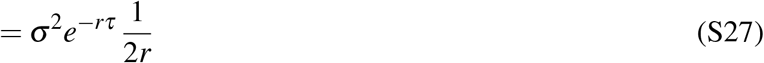

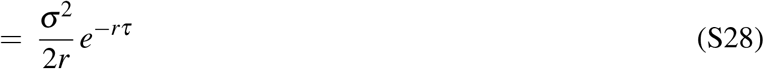

And for lag *τ* the autocovariance is:

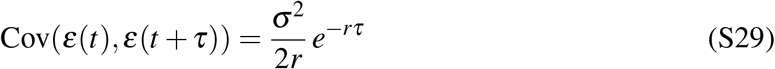

Thus, the population autocorrelation (ACF_*X*_ (*τ*)) for lag *τ* decays exponentially as:

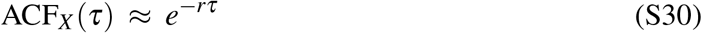

As discussed in the main manuscript, the OU model captures the exponential decay of autocorrelation. Ecologically, the parameter *r* thus serves as the relaxation rateit quantifies how quickly perturbations away from *K* are damped. A larger *r* means that deviations decay more rapidly (short memory of shocks), whereas a smaller *r* means that disturbances persist longer in time. Thus, the relaxation rate measures the resilience of the population to stochastic perturbations. We calculated the empirical autocorrelation function with and without seasonal effects, and show guppy populations should exhibit OU-like relaxation with decay rate ≈ *r*.

Also, note that with discrete observations every Δ*t*, the OU reduces to an autoregressive AR(1) process with autocorrelation *ρ*_*k*_ = *e*^−*rk*Δ*t*^.

### Appendix S4 Power spectral density and temporal stability

Population fluctuations can be studied in the time domain (via autocorrelation function or in the frequency domain (via power spectral density). The two are equivalent by the Wiener-Khinchin theorem: for a stationary process, the power spectral density is the Fourier transform of the auto-covariance function [5]. Thus, temporal stability can equivalently be characterized by the decay rate of the autocorrelation function (relaxation rate *r* in our OU approximation) or by the concentration of spectral power at low frequency [6].

For population deviations *ε*(*t*), power spectral density (Φ(*ω*)) is evaluated as:

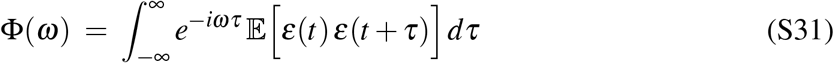

We know the autocovariance from equation (S29) is:

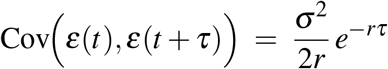

Thus, the power spectral density is can be evaluated as follows:

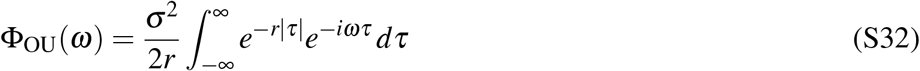

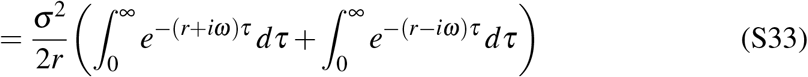

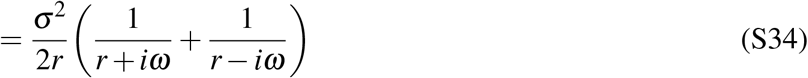

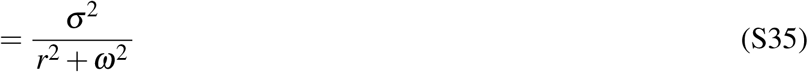

In our case, *σ* = *β K*,

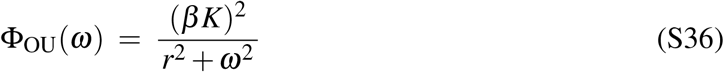

This spectrum describes how variance is distributed across temporal frequencies *ω* (cycles per unit time). This is the frequency-domain counterpart of the exponential ACF *ρ*(*τ*) = *e*^−*rτ*^ discussed above. The spectrum has a simple shape: it is highest at low frequencies (long timescales) and declines smoothly as frequency increases. Intuitively, this means most of the variance comes from slow fluctuations, while high-frequency oscillations are strongly damped. The rate of decline is set by the relaxation rate *r*: larger *r* shifts variance toward faster timescales (higher frequencies and faster decay of perturbations), while smaller *r* concentrates variance at long timescales (low frequency and longer memory of perturbations).

#### S4.1 From observed to environmental fluctuations

So far we considered the OU process driven by white noise. More generally, after linearization the deviations *ε* satisfy a first-order linear equation

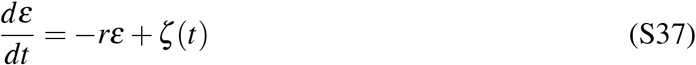

where *ζ* (*t*) represents stochastic environmental inputs. We assume this is a stable linear time-invariant system and *r* > 0. A standard way to characterize such systems is to ask: how do they respond to a brief pulse *ζ* (*t*) = δ (*t*)? Here, δ is the Dirac delta function and represents an instantaneous unit “kick” applied at *t* = 0. Substituting gives

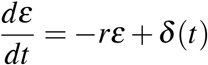

whose solution is zero for *t* < 0, jumps at *t* = 0, and then decays exponentially according to 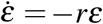. The resulting impulse response is

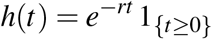

an exponentially decaying trajectory at rate *r*. By linearity and time-invariance, the response to the pulse inputs is a superposition of impulse responses. This yields the convolution representation

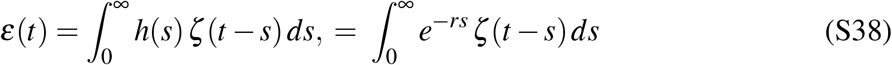

which shows that the present deviation is a weighted average of past inputs, which decay exponentially at rate *r*. Each input impulse generates its own exponential response, and the total state *ε*(*t*) is the sum of these overlapping decays.

A more convenient view of the convolution form in equation (S38) comes from the frequency domain: convolution in time corresponds to multiplication in frequency. The Fourier transform of the impulse response *h*(*t*) is the transfer function

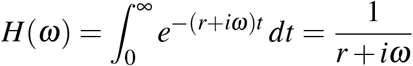

whose squared magnitude

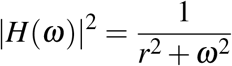

quantifies how inputs at frequency *ω* are transmitted to the state. Now let Φ(*ω*) be the spectral density of the output and *S*_*ζ*_ (*ω*) be the input power spectral density. In signal-processing [5], the state and input power spectral densities are linked by the relation:

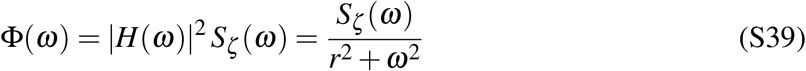

where Φ(*ω*) is the state spectrum and *S*_*ζ*_ (*ω*) is the input spectrum. Inverting (S39) gives the implied input spectrum:

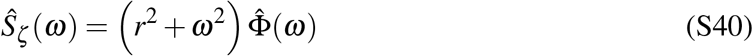

Given the observed state spectrum 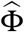and an estimate of the relaxation rate *r*, we can infer the frequency structure of the effective environmental drivers 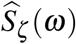 (up to a constant) from the observed population spectrum. Note that the inverse does not recover the true spectrum uniquely, but rather the effective spectrum consistent with the observed dynamics under the linearized OU approximation. If the driving noise is spectrally white (fluctuations have equal power at all frequencies), then 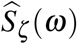 is expected to be constant across frequencies. Deviations from flatness therefore signal dominant extrenal timescales.

### Appendix S5 EWS metrics

We computed standard early warning signal (EWS) metrics for all the four stream populations. The metrics include lag-1 autocorrelation and variance using rolling window estimates discussed in Scheffer et al. [7] and Dakos et al. [8]. The idea is that systems show similar behaviour when they approach a critical transition: their variance and autocorrelation increase. This is because near a bifurcation point, the recovery rate of the system progressively deteriorates. So tipping points can be detected by indicators, such as the variance, temporal/spatial correlation, and recovery rate (recently summarized in Hastings et al. [9].

For LL stream, the population exhibits relatively stable dynamics with moderate fluctuations. The AR(1) signal shows a slight decline over time rather than the expected increase under critical slowing down. Variance increases gradually over part of the time series but then decreases. The population may be stabilizing following recent decline. For CA while variance exhibits a gradual increase, the AR(1) signal does not show a consistent upward trend indicative of critical slowing down.

FOr TY, the population shows larger fluctuations and higher variability compared to intact canopy streams. The AR(1) metric remains relatively flat with transient deviations but no sustained upward trend. Variance increases over time before declining toward the end of the series. These mixed patterns indicate no significant trends in EWS metrics. For UL, the population also exhibits large fluctuations and variability. The AR(1) signal shows non-monotonic behavior, including periods of decline and recovery, without a consistent increase. Variance increases over time and then stabilizes or declines slightly. There is no significant trend in EWS metrics.

**Figure S.1:**
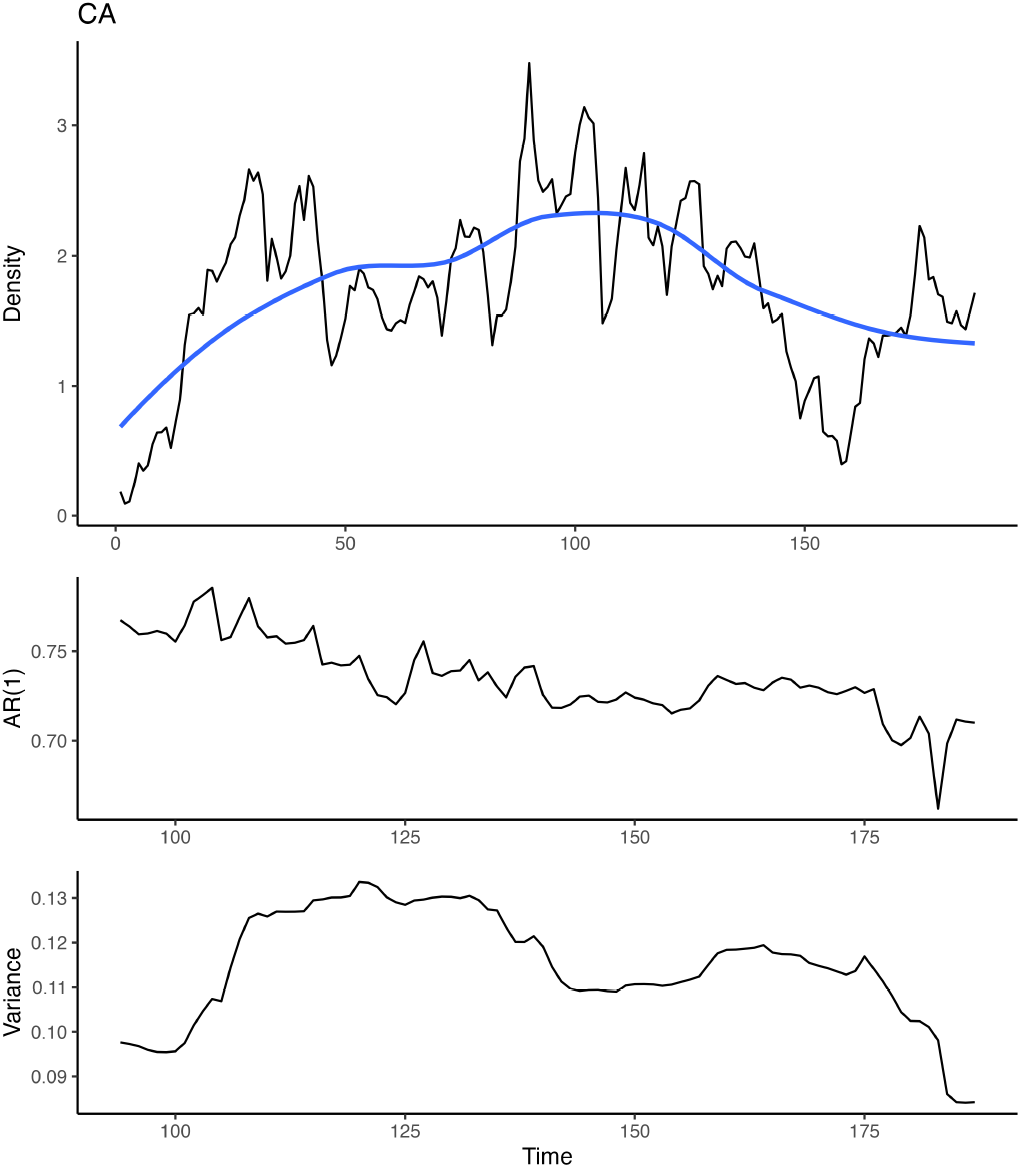
Early warning signal metrics for the LL population. Top panel shows the observed population density time series (black) with a Gaussian detrended smooth (blue). Middle panel shows the lag-1 autocorrelation (AR(1)) estimated over rolling windows. Bottom panel shows the corresponding variance. Time is measured in months. No consistent increase in AR(1) is observed, while variance shows a gradual increase over time.

**Figure S.2:**
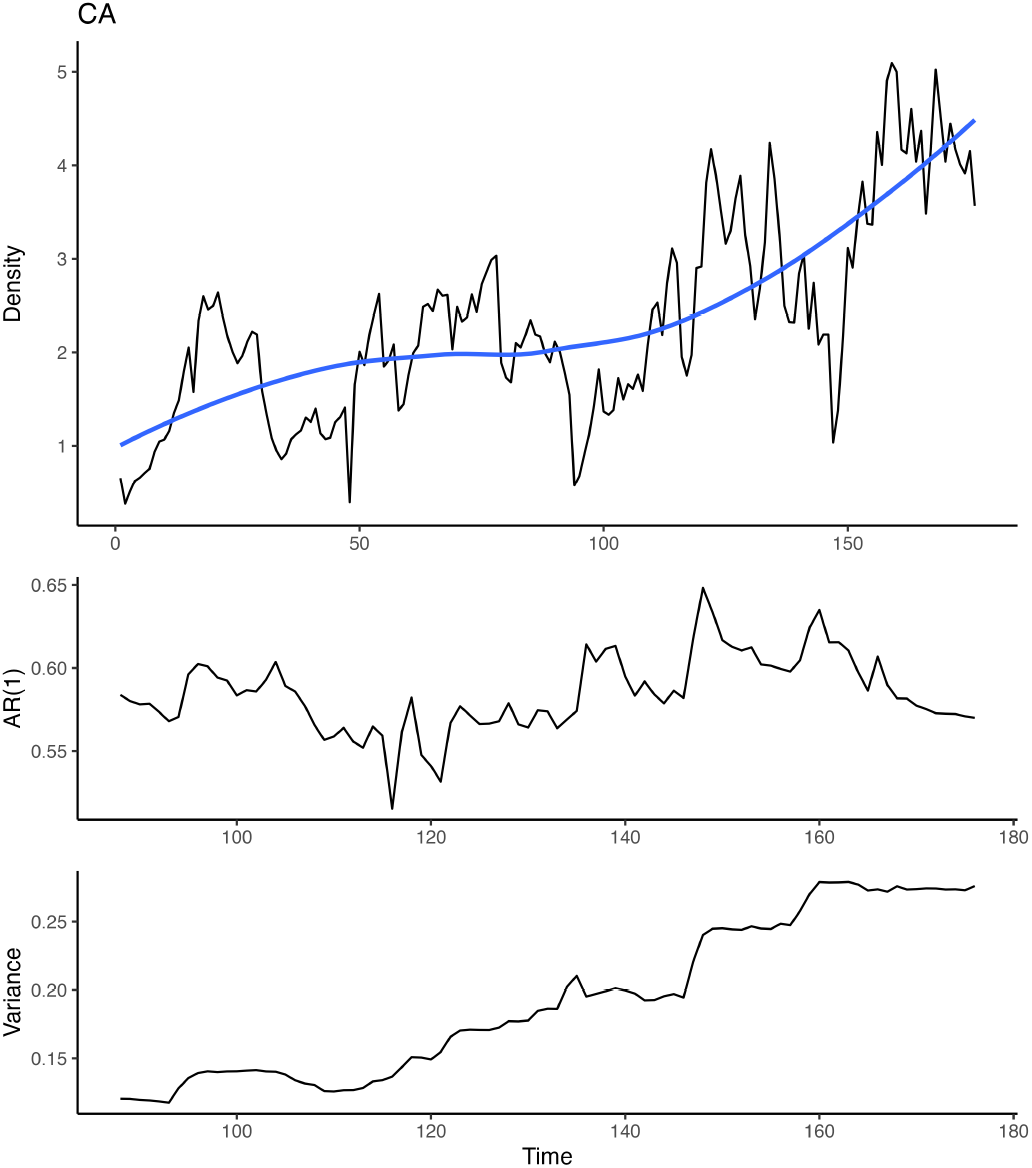
Early warning signal metrics for the CA population. Top panel shows the observed population density time series (black) with a Gaussian detrended smooth (blue). Middle panel shows the lag-1 autocorrelation (AR(1)) estimated over rolling windows. Bottom panel shows the corresponding variance. Time is measured in months. No consistent increase in AR(1) is observed, while variance shows a gradual increase over time.

**Figure S.3:**
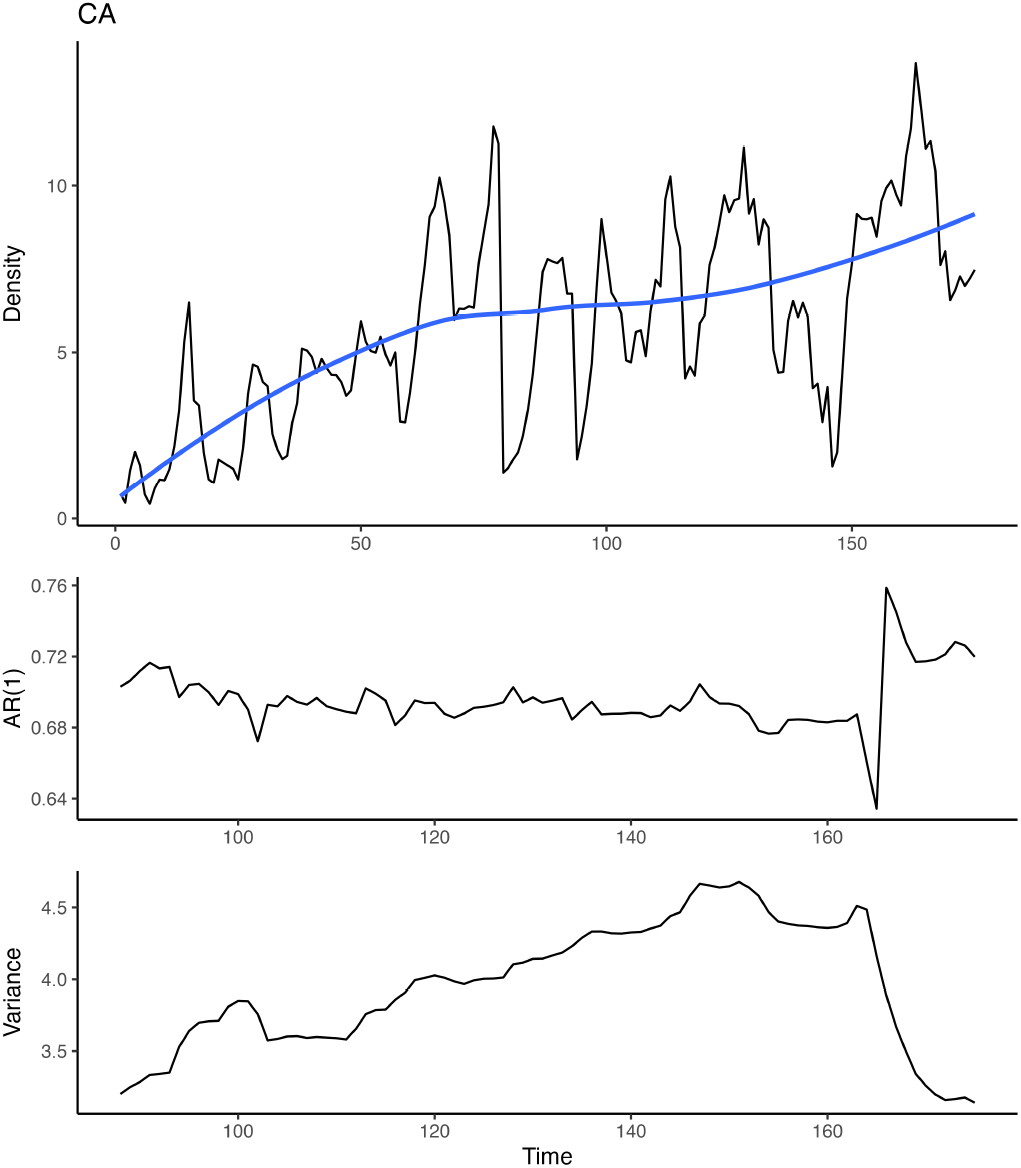
Early warning signal metrics for the TY population. Top panel shows the observed population density time series (black) with a Gaussian detrended smooth (blue). Middle panel shows the lag-1 autocorrelation (AR(1)) estimated over rolling windows. Bottom panel shows the corresponding variance. Time is measured in months. No consistent increase in AR(1) is observed, while variance shows a gradual increase over time.

**Figure S.4:**
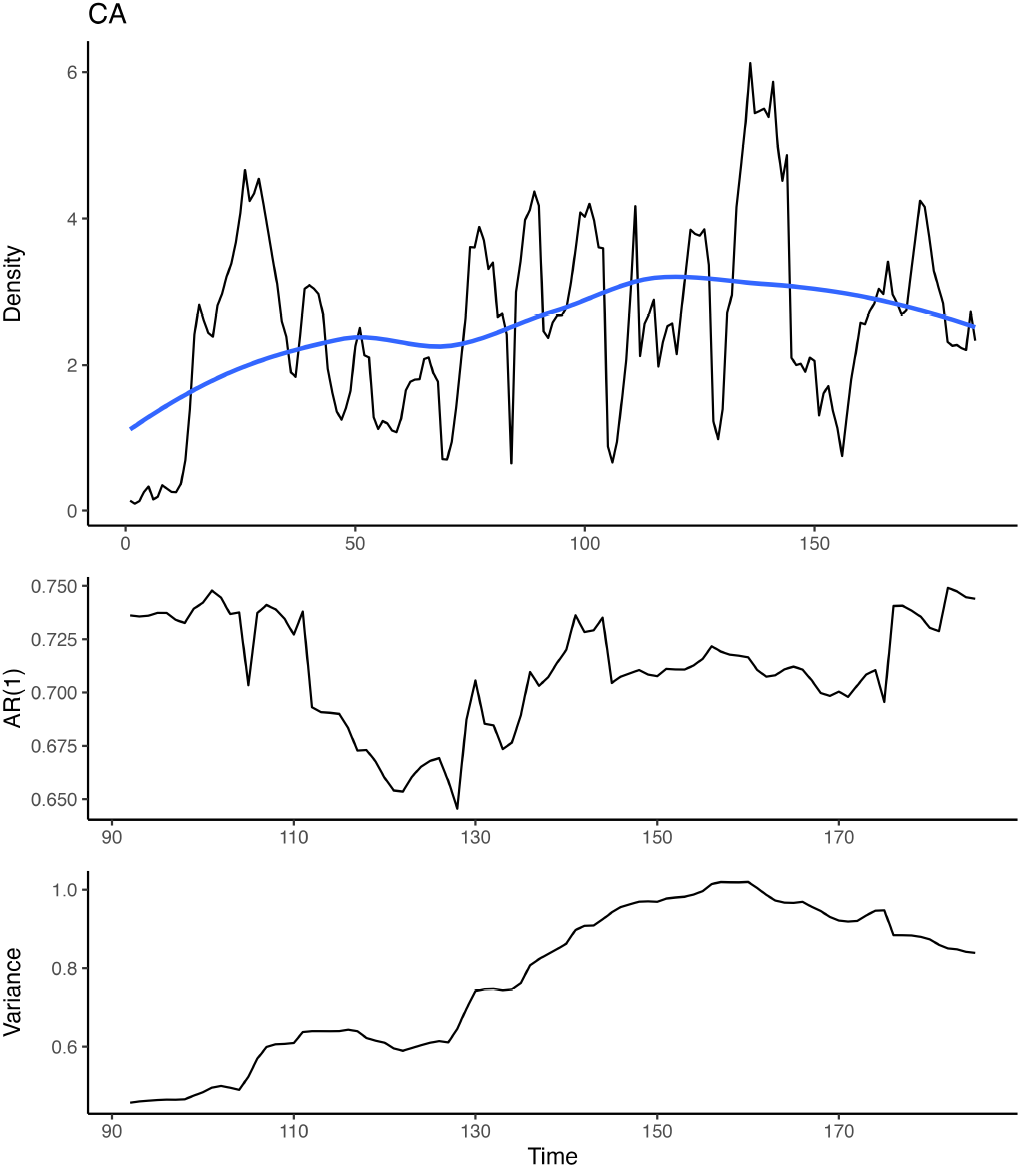
Early warning signal metrics for the UL population. Top panel shows the observed population density time series (black) with a Gaussian detrended smooth (blue). Middle panel shows the lag-1 autocorrelation (AR(1)) estimated over rolling windows. Bottom panel shows the corresponding variance. Time is measured in months. No consistent increase in AR(1) is observed, while variance shows a gradual increase over time.

**Figure S.5:**
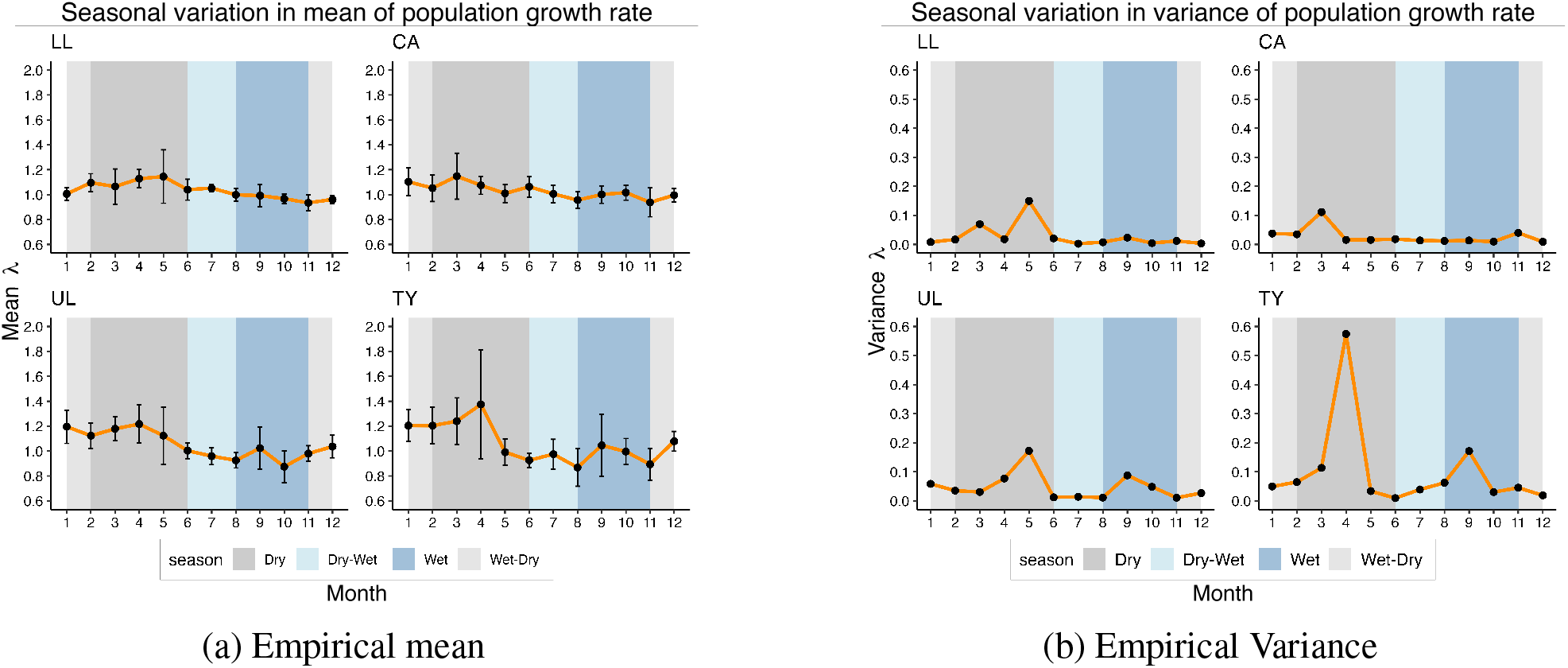
Empirical mean and variance across seasons.

### Appendix S6 Life-history correlates

We examine the life-hisotry correlates for mean and variance of population growth rate. Both empirical and estimated mean and variance of population growth rate (*λ* ) are highest in the transition from the dry season to the wet season (Figures S.5a and S.5b). These periods also correspond to the highest population densities due to increased recruitment and survival in the dry season. Given the state-dependent nature of fluctuations observed in our SDE analysis results, we hypothesize that seasonal peaks in *λ* variability may be driven by increased population densities during these times, which is itself driven by seasonality. Both TY and UL streams show higher increase in variance and mean growth rate (during the dry-wet transition) compared to the intact canopy streams LL and CA. Unlike other streams, TY also shows a negative correlation between survival and recruitment during the dry-wet months as shown in Fig S.5.

#### S6.1 Mean-Variance scaling

We further examined the scaling between the mean and variance of monthly population density by computing the mean and variance across months (for each stream and year) and fitting a log-log relationship between annual variance and mean (Taylor’s law). In the raw data, all streams exhibit positive mean-variance scaling, although the exponent varies across streams (Fig S.6).

To assess the role of seasonality, we removed the mean seasonal cycle by subtracting the long-term monthly mean (computed across years for each stream) and adding back the overall mean. After deseasonalization, this relationship weakens substantially in LL, UL, and TY, whereas the intact canopy stream (CA) retains a positive scaling relationship (Fig S.6).

These patterns are consistent with the spectral and autocorrelation analyses, indicating that mean-variance scaling in most streams is largely driven by seasonal fluctuations, whereas in CA it reflects intrinsic population variability.

**Figure S.6:**
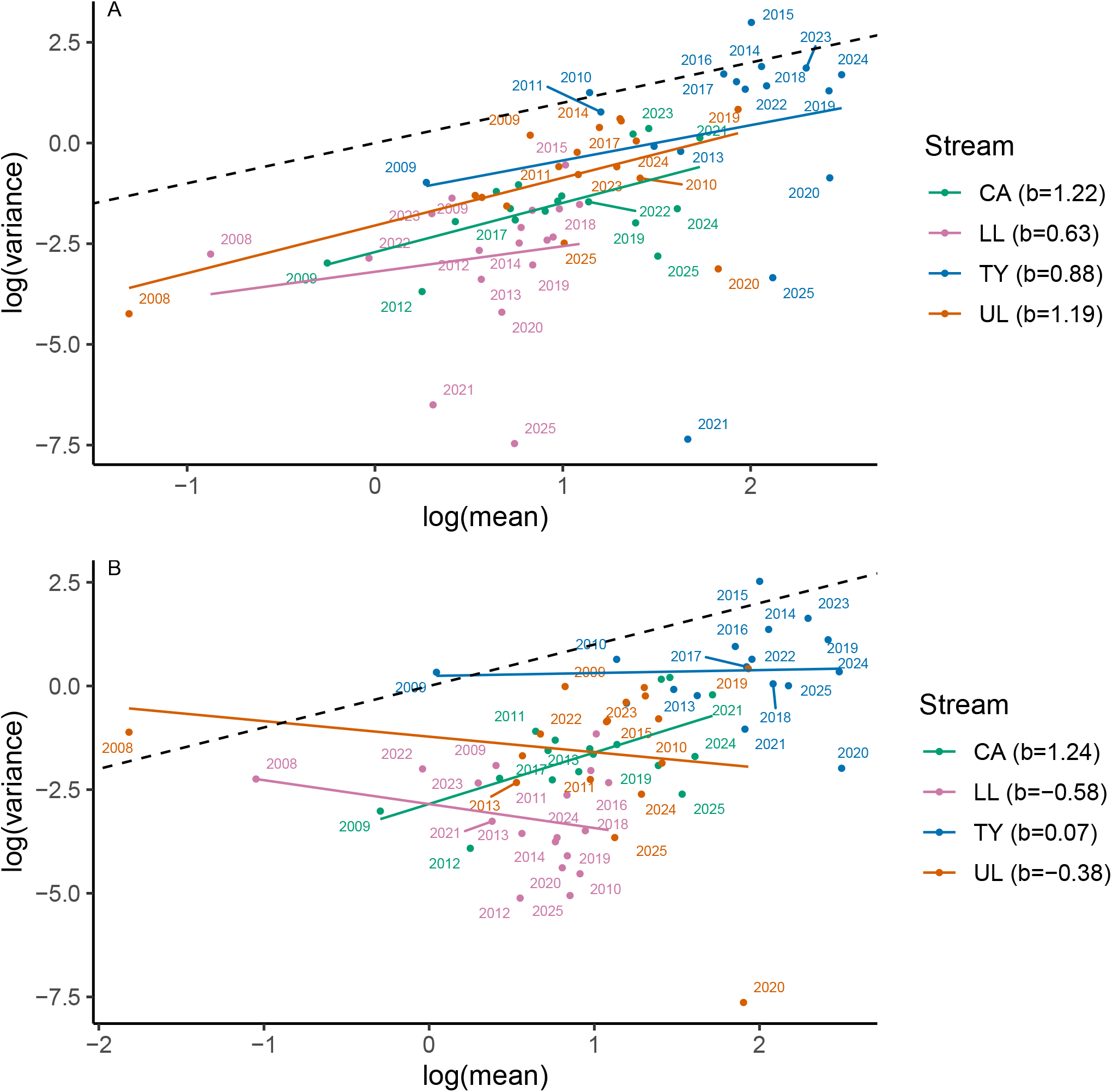
Relationship between mean and variance of population density on log-log scale. The top panel (A) depicts the relaitonship computed for raw population densities (computed from monthly observations within each year) for each stream. Points represent years and colored lines denote linear fits for each streams. The slopes are shown in the legend. The dashed line indicates the reference slope b=1. The bottom panel (B) shows the same but after removing the mean seasonal cycle by subtracting the long-term monthly mean and adding back the overall mean for each stream.

### Appendix S7 Validation of the equation discovery procedure

Supplement figure S.7 shows the model and noise-diagnostic tests for the equation discovery method, as outlined in [1]. The model diagnostics compare the histogram and autocorrelation function of the observed data with the theoretical expectations, namely a Gamma distribution and an exponential decay respectively (Appendix S2, S3). For all streams, we see good agreement between the data and the theoretical expectation based on the discovered model, barring the seasonal fluctuations found in the real data. The noise diagnostic tests look at the distribution and autocorrelation of the residuals, estimated as 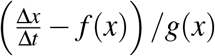. The distribution of the residuals match a standard Gaussian distribution for all streams. The residual autocorrelation functions also decay quickly (barring seasonal fluctuations which are unaccounted for by the logistic model), validating the Markovian assumptions.

Taken together, these diagnostic tests show that the population dynamics in all streams are well-approximated by the stochastic logistic equations discovered by the equation-discovery method, with the residual fluctuations accounted for by seasonal effects (Section 2.4).

**Figure S.7:**
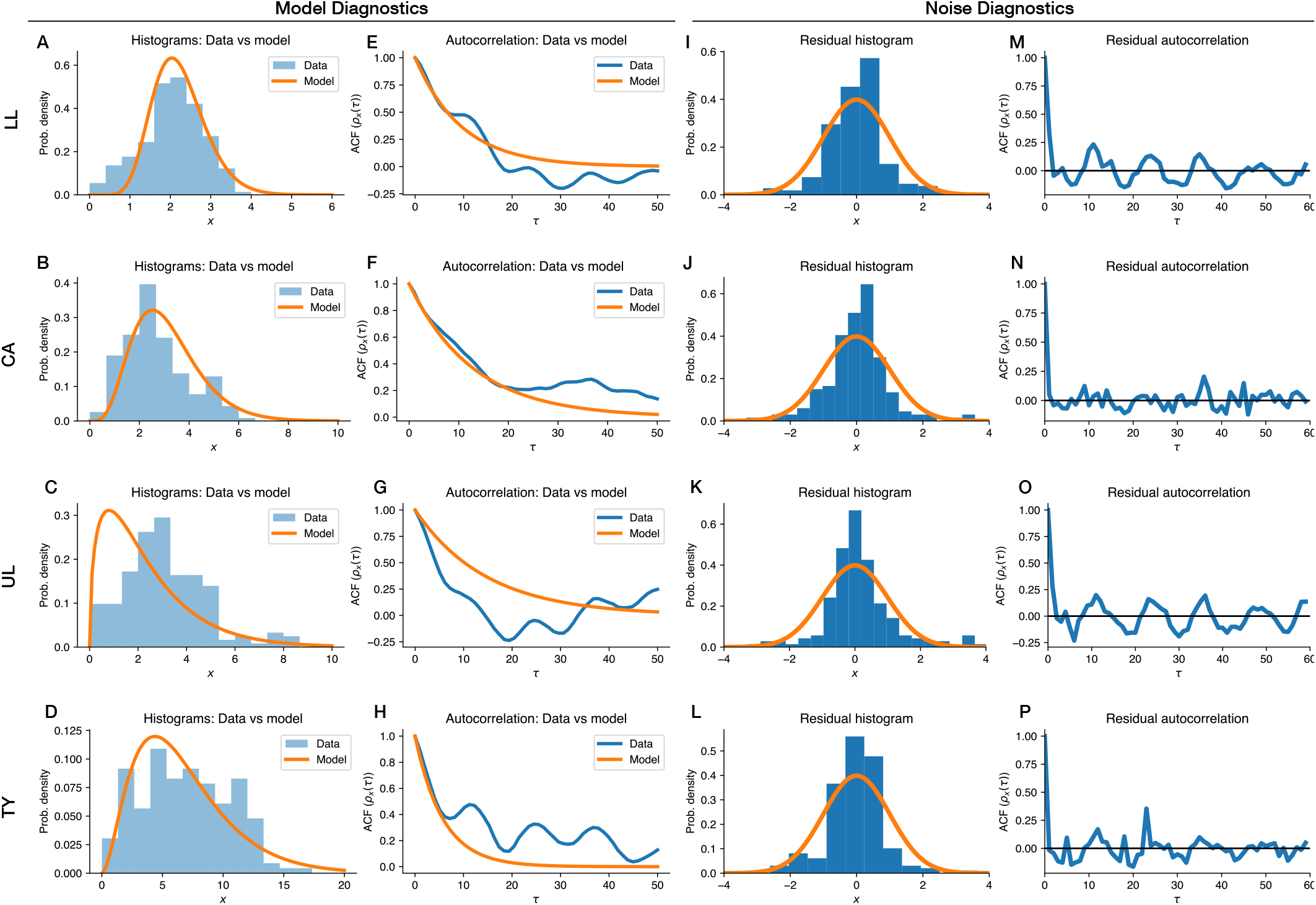
Diagnostics for the discovered SDE models. (A-D) Comparison of the data histograms (blue) with the theoretically expected distribution based on the model (orange lines). (E-H) Comparison of the autocorrelation function of the data with the theoretical expectation. (I-L) Comparison of the histogram of the residual fluctuations with a theoretically expected standard normal distribution (M-P) Autocorrelation functions of the residual fluctuations.

## References

[1] Karen Filbee-Dexter and Robert E Scheibling. “Sea urchin barrens as alternative stable states of collapsed kelp ecosystems”. Marine ecology progress series 495 (2014), pp. 1–25.

[2] David Alonso, Alan J McKane, and Mercedes Pascual. “Stochastic amplification in epidemics”. Journal of the Royal Society Interface 4.14 (2007), pp. 575–582.

[3] Russell Lande. “Risks of population extinction from demographic and environmental stochasticity and random catastrophes”. The American Naturalist 142.6 (1993), pp. 911–927.

[4] Brett A Melbourne and Alan Hastings. “Extinction risk depends strongly on factors contributing to stochasticity”. Nature 454.7200 (2008), pp. 100–103.

[5] Joseph Travis et al. “Density-dependent selection”. Annual Review of Ecology, Evolution, and Systematics 54.1 (2023), pp. 85–105.

[6] Manvi Sharma et al. “Can livestock grazing dampen density-dependent fluctuations in wild herbivore populations?” Journal of Applied Ecology 61.6 (2024), pp. 1243–1254.

[7] Xavier Lambin et al. “Density-dependent recruitment but not survival drives cyclic dynamics in a field vole population”. Proceedings of the National Academy of Sciences 122.40 (2025), e2509516122.

[8] Arpat Ozgul et al. “The dynamics of phenotypic change and the shrinking sheep of St. Kilda”. Science 325.5939 (2009), pp. 464–467.

[9] Isabel A Honda et al. “Seasonal trophic controls drive population variability in a foundational marine copepod”. Scientific Reports 15.1 (2025), p. 36018.

[10] Shripad Tuljapurkar, Jean-Michel Gaillard, and Tim Coulson. “From stochastic environments to life histories and back”. Philosophical Transactions of the Royal Society B: Biological Sciences 364.1523 (2009), pp. 1499–1509.

[11] Robin E Snyder and Stephen P Ellner. “Pluck or luck: does trait variation or chance drive variation in lifetime reproductive success?” The American Naturalist 191.4 (2018), E90–E107.

[12] Peter R Grant, B Rosemary Grant, Lukas F Keller, and Kenneth Petren. “Effects of El Niño events on Darwin’s finch productivity”. Ecology 81.9 (2000), pp. 2442–2457.

[13] Vadim A Karatayev, Stephan B Munch, Tanya L Rogers, and Daniel C Reuman. “Climate change could amplify weak synchrony in large marine ecosystems”. Proceedings of the National Academy of Sciences 122.1 (2025), e2404155121.

[14] Russell Lande, Steinar Engen, and Bernt-Erik Sæther. “An evolutionary maximum principle for density-dependent population dynamics in a fluctuating environment”. Philosophical Transactions of the Royal Society B: Biological Sciences 364.1523 (2009), pp. 1511–1518.

[15] Peter Turchin. “Nonlinear time-series modeling of vole population fluctuations”. Population Ecology 38.2 (1996), pp. 121–132.

[16] BT Grenfell et al. “Noise and determinism in synchronized sheep dynamics”. Nature 394.6694 (1998), pp. 674–677.

[17] Jean-Michel Gaillard and Nigel Gilles Yoccoz. “What does drive temporal variation in population size in mammalian species?” Proceedings of the National Academy of Sciences 122.49 (2025), e2526456122.

[18] N Chr Stenseth et al. “Dynamics of coastal cod populations: intra-and intercohort density dependence and stochastic processes”. Proceedings of the Royal Society of London. Series B: Biological Sciences 266.1429 (1999), pp. 1645–1654.

[19] Brian Dennis and Mark RM Otten. “Joint effects of density dependence and rainfall on abundance of San Joaquin kit fox”. The Journal of wildlife management (2000), pp. 388–400.

[20] Russell Lande, Steinar Engen, and Bernt-Erik Saether. Stochastic population dynamics in ecology and conservation. OUP Oxford, 2003.

[21] Bernt-Erik Sæther et al. “Demographic routes to variability and regulation in bird populations”. Nature communications 7.1 (2016), p. 12001.

[22] Brian Dennis and Ganapati P Patil. “The gamma distribution and weighted multimodal gamma distributions as models of population abundance”. Mathematical Biosciences 68.2 (1984), pp. 187–212.

[23] Steinar Engen and Russell Lande. “Population Dynamic Models Generating Species Abundance Distributions of the Gamma Type”. en. Journal of Theoretical Biology 178.3 (Feb. 1996), pp. 325–331. ISSN: 00225193. DOI: 10.1006/jtbi.1996.0028. URL: https://linkinghub.elsevier.com/retrieve/pii/S0022519396900284 (visited on 12/20/2025).

[24] Paolo D’Odorico, Francesco Laio, and Luca Ridolfi. “Noise-induced stability in dryland plant ecosystems”. Proceedings of the National Academy of Sciences 102.31 (2005), pp. 10819–10822.

[25] Vishwesha Guttal and C Jayaprakash. “Impact of noise on bistable ecological systems”. Ecological modelling 201.3-4 (2007), pp. 420–428.

[26] Carl Boettiger. “From noise to knowledge: how randomness generates novel phenomena and reveals information”. Ecology letters 21.8 (2018), pp. 1255–1267.

[27] Hongxia Zhang, Wei Xu, Youming Lei, and Yan Qiao. “Noise-induced vegetation transitions in the grazing ecosystem”. Applied Mathematical Modelling 76 (2019), pp. 225–237.

[28] Jitesh Jhawar et al. “Noise-induced schooling of fish”. Nature Physics 16.4 (2020), pp. 488–493.

[29] Ludwig Arnold. “Random dynamical systems”. Dynamical Systems: Lectures Given at the 2nd Session of the Centro Internazionale Matematico Estivo (CIME) held in Montecatini Terme, Italy, June 13–22, 1994. Springer, 2006, pp. 1–43.

[30] Marc Mendler, Johannes Falk, and Barbara Drossel. “Analysis of stochastic bifurcations with phase portraits”. PloS one 13.4 (2018), e0196126.

[31] Janez Gradišek, Silke Siegert, Rudolf Friedrich, and Igor Grabec. “Analysis of time series from stochastic processes”. Physical Review E 62.3 (2000), p. 3146.

[32] Rahimi Tabar. Analysis and data-based reconstruction of complex nonlinear dynamical systems. Vol. 730. Springer, 2019.

[33] Arshed Nabeel et al. “Discovering stochastic dynamical equations from ecological time series data”. The American Naturalist 205.4 (2025), E100–E117.

[34] Vasilis Dakos et al. “Methods for detecting early warnings of critical transitions in time series illustrated using simulated ecological data”. PloS one 7.7 (2012), e41010.

[35] Sarah J Burthe et al. “Do early warning indicators consistently predict nonlinear change in long-term ecological data?” Journal of Applied Ecology 53.3 (2016), pp. 666–676.

[36] Li Xu, Denis Patterson, Simon Asher Levin, and Jin Wang. “Non-equilibrium early-warning signals for critical transitions in ecological systems”. Proceedings of the National Academy of Sciences 120.5 (2023), e2218663120.

[37] Alan Hastings, Sergei Petrovskii, Valerio Lucarini, and Andrew Morozov. “Tipping points in complex ecological systems”. arXiv preprint arXiv:2602.20702 (2026).

[38] Alison J Robey and David A Vasseur. “Order matters: Autocorrelation of temperature dictates extinction risk in populations with nonlinear thermal performance”. Ecology 107.3 (2026), e70325.

[39] Shripad Tuljapurkar and CV Haridas. “Temporal autocorrelation and stochastic population growth”. Ecology letters 9.3 (2006), pp. 327–337.

[40] Harman Jaggi et al. “Transient dynamics and nonlinear fitness: a unified matrix approach to press and pulse perturbation”. BioRxiv (2025), pp. 2025–09.

[41] Maurice Bertram Priestley. The spectral analysis of time series. 1988.

[42] Yvonne Krumbeck, Qian Yang, George WA Constable, and Tim Rogers. “Fluctuation spectra of large random dynamical systems reveal hidden structure in ecological networks”. Nature Communications 12.1 (2021), p. 3625.

[43] Sarah M Collins et al. “Fish introductions and light modulate food web fluxes in tropical streams: a whole-ecosystem experimental approach”. Ecology 97.11 (2016), pp. 3154–3166.

[44] Antoine OHC Leduc et al. “The experimental range extension of guppies (Poecilia reticulata) influences the metabolic activity of tropical streams”. Oecologia 195.4 (2021), pp. 1053–1069.

[45] Tyler J Kohler et al. “Flow, nutrients, and light availability influence Neotropical epilithon biomass and stoichiometry”. Freshwater Science 31.4 (2012), pp. 1019–1034.

[46] David A Reznick, Heather Bryga, and John A Endler. “Experimentally induced life-history evolution in a natural population”. Nature 346.6282 (1990), pp. 357–359.

[47] Ronald D Bassar et al. “Local adaptation in Trinidadian guppies alters ecosystem processes”. Proceedings of the National Academy of Sciences 107.8 (2010), pp. 3616–3621.

[48] Joseph Travis et al. “Population regulation and density-dependent demography in the Trinidadian guppy”. The American Naturalist 202.4 (2023), pp. 413–432.

[49] Zhuan Cheng, Jinqiao Duan, and Liang Wang. “Most probable dynamics of some nonlinear systems under noisy fluctuations”. Communications in Nonlinear Science and Numerical Simulation 30.1-3 (2016), pp. 108–114.

[50] JH Yang et al. “Stochastic P-bifurcation and stochastic resonance in a noisy bistable fractionalorder system”. Communications in Nonlinear Science and Numerical Simulation 41 (2016), pp. 104–117.

[51] Ping Han et al. “The stochastic P-bifurcation analysis of the impact system via the most probable response”. Chaos, Solitons & Fractals 144 (2021), p. 110631.

[52] SR Carpenter and WA Brock. “Early warnings of unknown nonlinear shifts: a nonparametric approach”. Ecology 92.12 (2011), pp. 2196–2201.

[53] Vasilis Dakos et al. “Tipping point detection and early warnings in climate, ecological, and human systems”. Earth System Dynamics 15.4 (2024), pp. 1117–1135.

[54] Richard C Lewontin and Daniel Cohen. “On population growth in a randomly varying environment”. Proceedings of the National Academy of sciences 62.4 (1969), pp. 1056–1060.

[55] Shripad Tuljapurkar and Wenyun Zuo. “Mutations and the distribution of lifetime reproductive success”. Journal of the Indian Institute of Science 102.4 (2022), pp. 1269–1275.

[56] Sha Jiang et al. “Reproductive dispersion and damping time scale with life-history speed”. Ecology Letters 25.9 (2022), pp. 1999–2008.

[57] Shripad D Tuljapurkar. “Population dynamics in variable environments. III. Evolutionary dynamics of r-selection”. Theoretical population biology 21.1 (1982), pp. 141–165.

[58] Russell Lande and Steven Hecht Orzack. “Extinction dynamics of age-structured populations in a fluctuating environment.” Proceedings of the National Academy of Sciences 85.19 (1988), pp. 7418–7421.

[59] Otso Ovaskainen and Baruch Meerson. “Stochastic models of population extinction”. Trends in ecology & evolution 25.11 (2010), pp. 643–652.

[60] Peter Chesson. “General theory of competitive coexistence in spatially-varying environments”. Theoretical population biology 58.3 (2000), pp. 211–237.

[61] Stephen P Ellner, Robin E Snyder, Peter B Adler, and Giles Hooker. “An expanded modern coexistence theory for empirical applications”. Ecology letters 22.1 (2019), pp. 3–18.

[62] David A Vasseur and Peter Yodzis. “The color of environmental noise”. Ecology 85.4 (2004), pp. 1146–1152.

[63] Shripad Tuljapurkar. Population dynamics in variable environments. Springer Science & Business Media, 2013.

[64] George Sugihara et al. “Empirical dynamic modeling”. The Comprehensive R Archive Network (2020).

[65] Stephan B Munch, Tanya L Rogers, and George Sugihara. “Recent developments in empirical dynamic modelling”. Methods in Ecology and Evolution 14.3 (2023), pp. 732–745.

[66] Ottar N Bjørnstad and Bryan T Grenfell. “Noisy clockwork: time series analysis of population fluctuations in animals”. Science 293.5530 (2001), pp. 638–643.

[67] Marten Scheffer et al. “Early-warning signals for critical transitions”. Nature 461.7260 (2009), pp. 53–59.

[68] Harman Jaggi et al. “Density dependence shapes life-history trade-offs in a food-limited population”. Ecology Letters 27.11 (2024), e14551.

[69] M Kajin et al. “Second-order elasticities for Ecology and Evolution: Unravelling nonlinear fitness responses to perturbations”. bioRxiv (2025), pp. 2025–04.

[70] Shripad Tuljapurkar, Harman Jaggi, and Wenyun Zuo. “Generalizing the Theory to Include Age and Stage” (2025).

[71] Ronald D Bassar, Andres Lopez-Sepulcre, David N Reznick, and Joseph Travis. “Experimental evidence for density-dependent regulation and selection on Trinidadian guppy life histories”. The American Naturalist 181.1 (2013), pp. 25–38.

[72] Christina Maria Hernandez et al. “A robust method for quantifying the contribution of transient dynamics to variation in population growth rate” (2026).

[73] Jacopo Grilli. “Macroecological laws describe variation and diversity in microbial communities”. Nature communications 11.1 (2020), p. 4743.

[74] Silvia Zaoli and Jacopo Grilli. “The stochastic logistic model with correlated carrying capacities reproduces beta-diversity metrics of microbial communities”. en. PLOS Computational Biology 18.4 (Apr. 2022). Ed. by Tobias Bollenbach, e1010043. ISSN: 1553-7358. DOI: 10.1371/journal.pcbi.1010043. URL: https://dx.plos.org/10.1371/journal.pcbi.1010043 (visited on 12/01/2025).

[75] Joseph Travis et al. “Do eco-evo feedbacks help us understand nature? Answers from studies of the Trinidadian guppy”. Advances in ecological research. Vol. 50. Elsevier, 2014, pp. 1–40.

[76] Sonya K Auer et al. “Life histories have a history: effects of past and present conditions on adult somatic growth rates in wild Trinidadian guppies”. Journal of Animal Ecology 81.4 (2012), pp. 818–826.

[77] Tomos Potter et al. “Environmental change, if unaccounted, prevents detection of cryptic evolution in a wild population”. The American Naturalist 197.1 (2021), pp. 29–46.

[78] Jeffrey Lee Laake. “RMark: an R interface for analysis of capture-recapture data with MARK” (2013).

[79] Werner Horsthemke and René Lefever. Noise-induced transitions: theory and applications in physics, chemistry, and biology. Springer, 1984.

[80] Crispin W Gardiner et al. Handbook of stochastic methods. Vol. 3. springer Berlin, 2004.

[81] John G Proakis. Digital signal processing: principles, algorithms, and applications, 4/E. Pearson Education India, 2007.

## Appendix References

[1] Arshed Nabeel et al. “Discovering stochastic dynamical equations from ecological time series data”. The American Naturalist 205.4 (2025), E100–E117.

[2] Marc Mendler, Johannes Falk, and Barbara Drossel. “Analysis of stochastic bifurcations with phase portraits”. PloS one 13.4 (2018), e0196126.

[3] H. Risken and H. Haken. The Fokker-Planck Equation: Methods of Solution and Applications Second Edition. Springer, 1989.

[4] Crispin W Gardiner et al. Handbook of stochastic methods. Vol. 3. springer Berlin, 2004.

[5] John G Proakis. Digital signal processing: principles, algorithms, and applications, 4/E. Pearson Education India, 2007.

[6] Yvonne Krumbeck, Qian Yang, George WA Constable, and Tim Rogers. “Fluctuation spectra of large random dynamical systems reveal hidden structure in ecological networks”. Nature Communications 12.1 (2021), p. 3625.

[7] Marten Scheffer et al. “Early-warning signals for critical transitions”. Nature 461.7260 (2009), pp. 53–59.

[8] Vasilis Dakos et al. “Methods for detecting early warnings of critical transitions in time series illustrated using simulated ecological data”. PloS one 7.7 (2012), e41010.

[9] Alan Hastings, Sergei Petrovskii, Valerio Lucarini, and Andrew Morozov. “Tipping points in complex ecological systems”. arXiv preprint arXiv:2602.20702 (2026).

